# Inferring fungal cis-regulatory networks from genome sequences via unsupervised and interpretable representation learning

**DOI:** 10.1101/2025.02.27.640643

**Authors:** Alan M Moses, Jason E Stajich, Audrey P Gasch, David A Knowles

## Abstract

Gene expression patterns are determined to a large extent by transcription factor binding to non-coding regulatory regions in the genome. However, gene expression cannot yet be systematically predicted from genome sequences, in part because non-functional matches to the sequence patterns (motifs) recognized by transcription factors (TFs) occur frequently throughout the genome. Large-scale functional genomics data for many TFs has enabled characterization of regulatory networks in experimentally accessible cells such as budding yeast. Beyond yeast, fungi are important industrial organisms and pathogens, but large-scale functional data is only sporadically available. Uncharacterized regulatory networks control key pathways and gene expression programs associated with fungal phenotypes. Here we explore a sequence-only approach to inferring regulatory networks by leveraging the 100s of genomes now available for many clades of fungi. We use gene orthology as the learning signal to infer interpretable, TF motif-based representations of non-coding regulatory regions. Using these representations to identify conserved signals for motifs, comparative genomics can be scaled to evolutionary comparisons where sequence similarity cannot be detected. We show that similarity of these conserved motif signals predicts gene expression and regulation better than using experimental data, and that we can infer known and novel regulatory connections in diverse fungi. Our new predictions include a pathway for recombination in *C. albicans* and pathways for mating and an RNAi immune response in *Neurospora*. Taken together, our results indicate that specific hypotheses about transcriptional regulation in fungi can be obtained for many genes from genome sequence analysis alone.

## Introduction

The ability to infer transcriptional regulatory networks from genome sequence alone is a long-standing goal of the post-genomic era: it would imply a basic understanding of the cis-regulatory code that connects regulatory sequences in the genome to gene expression patterns[1,2].

Although sequence-specific transcription factors (TFs) are appreciated as key determinants of transcription initiation [3], applying TF motif-discovery and motif-scanning to non-coding regions of single genomes is plagued by false positives [4,5]. It is long appreciated that evolutionary conservation can be used to increase the signal-to-noise because functional regulatory elements are more likely to be conserved [6–8]. Indeed, analyses of whole genome alignments have identified catalogues of TF binding motifs [9–11] and in some cases have power to identify individual regulatory connections between TFs and their targets[12].

In many clades, including the fungi considered here, regulatory regions diverge at the primary sequence level, even when functional information is conserved[13–16]. This means that functionally conserved non-coding regions may be difficult to align accurately beyond the most closely related species[16–18], so that alignment-based approaches to improving the signal-signal-to-noise of TF motif-discovery and scanning are limited. Several approaches have been developed to address this issue [17,19–22], but they do not readily scale to the large numbers of genomes currently available[23–25]. Fungal genomes represent a feasible case for regulatory network inference from sequence alone: they contain relatively short intergenic regions and introns, so automatic gene annotation pipelines are accurate, and most genes do not have multiple enhancers[26,27]. Predicting regulatory networks from sequence would have far-ranging applications. For diverse fungi, it is often not practical to perform systematic experiments because aspects of the life cycle are inaccessible in the lab[28]. Furthermore, for human and agriculture pathogens, lab experimental approaches are limited at least by regulations and at worst by dangers[29].

We frame inference of cis-regulatory networks from genome sequence as a representation learning problem[30]. Instead of using the DNA sequences directly, we aim to find a compact, quantitative and biologically interpretable representation for non-coding sequences[31] that can be used to understand their regulatory function. Using inference methods developed for deep learning, recent work aims to build mechanistic biophysical or bioinformatic models of transcriptional regulation by learning to predict large-scale experimental data [32–35]. Following this approach, we learn 256 TF binding motifs (position weight matrices, PWMs[36]) to represent orthologous sets of promoters.

Our key innovation is to learn a motif-based representation of regulatory regions from genome sequences alone. To do so, we infer PWMs and other parameters using reverse homology, a contrastive approach that maximizes evolutionary conservation in the representation space by leveraging shared ancestry as the learning signal[37,38]. Because our approach requires no direct experimental data, it can be applied to conditions and organisms for which large-scale functional genomics data is not available. In not relying on experimental data, our approach is similar to systematic phylogenetic footprinting[39] or unsupervised training of large-language-models[40] where motifs and regulatory connections can in some cases be identified through post-hoc analysis [41]. Unlike those approaches, however, our approach yields an interpretable motif-based representation of non-coding sequences from which regulatory connections can be read directly.

We find that patterns in the motif-based representations are strongly associated with gene expression and biological function, and that we can identify known and novel regulatory connections from visualizations of the representation space. Using this approach, we identify a putative recombination/parasex pathway in *C. albicans* and predict mating and virus-response pathways in *N. crassa*. Finally, we show that for the relatively unstudied genome of the pathogen *F. graminearum*, the promoter representation reveals a useful regulatory map, including predictions of regulatory mechanisms for effector genes. Our work demonstrates, in diverse fungi, the feasibility of scaling sequence-based regulatory network inference beyond alignable genomes, and yields rich biology, comparable to functional genomics approaches.

## Results

### Learning motif-based representations of orthologous sets of promoter regions without sequence alignments

At evolutionary distances where we can still identify orthologs for most genes, fungal non-coding regions upstream of the start codon (which we refer to subsequently as promoter regions, see Discussion) show conservation of regulatory function, but little sequence similarity. For example, the upstream regions of *S. cerevisiae PHO84* and its orthologs contain many matches to the Pho4 consensus site (Figure 1a), but these are only alignable in very closely related species (Figure 1a, Saccharomyces *sensu stricto*). The presence of motif matches in many orthologous promoters improves the signal-to-noise of motif-scanning and motif-discovery [10,19,22,31] because many matches are less likely to occur by chance. We sought to design an approach to leverage this improved signal-to-noise available in multiple orthologous promoter regions, without relying on sequence alignments.

**Figure 1.**
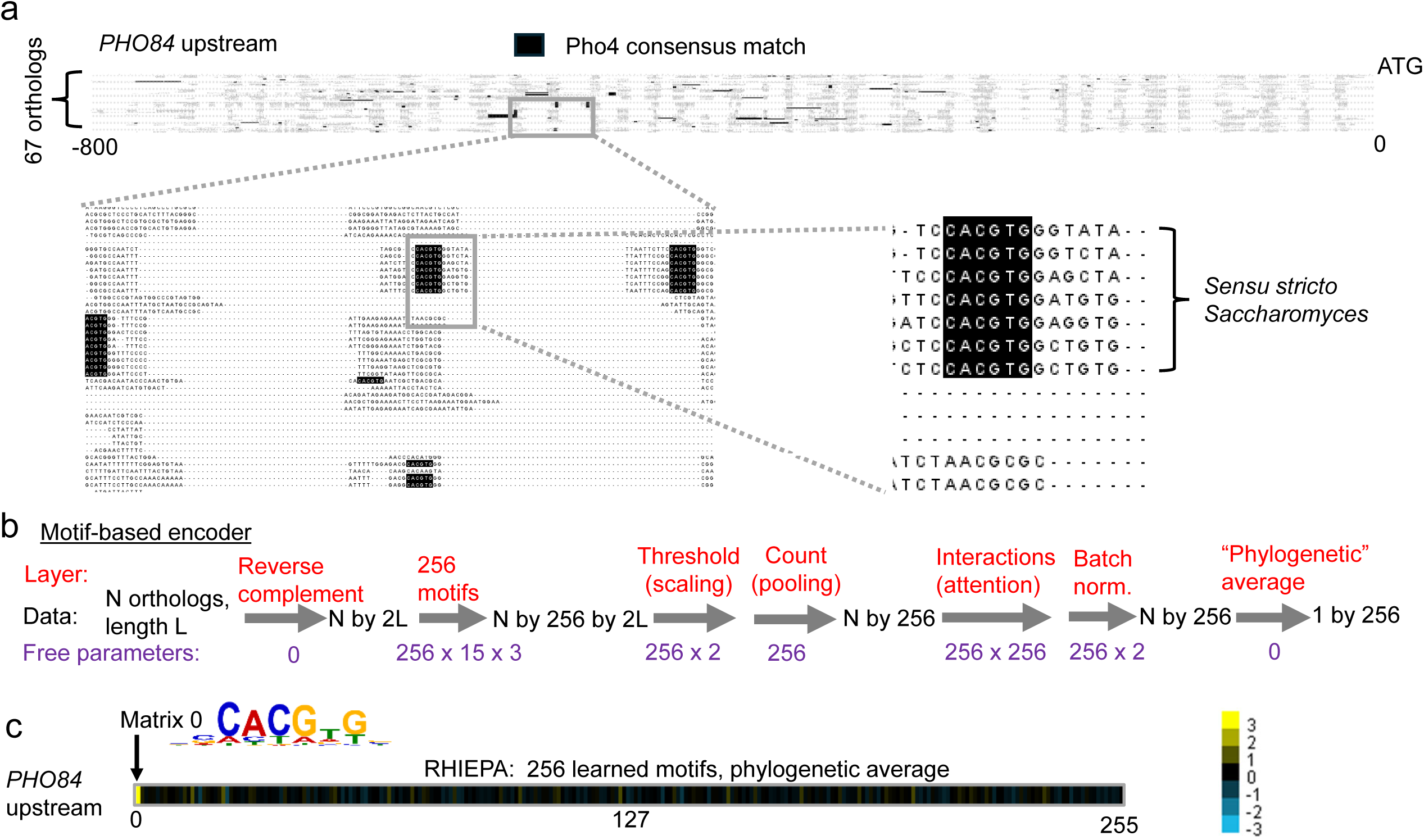
Motif-based encoder to represent orthologous sets of upstream regions. a) Shows the alignment of the upstream region for a single yeast promoter (*PHO84*) and orthologs. In yeast, *PHO84* is known to be a target of Pho4, and consensus sites for that transcription factor are found in the promoters from all species (black boxes), but are only aligned among the most closely related species (black box over text) b) a motif-based encoder changes the sequence data for an upstream region and its orthologs into a 256 dimensional numerical representation. Dimensions of data inputs and outputs are shown as black text, transformations (implemented as layers in Keras) are showed in red text and the number of free parameters in each layer is indicated in purple text. c) the reverse <u>h</u>omology, interpretable <u>e</u>ncoder, <u>p</u>hylogenetic <u>a</u>verage (RHIEPA) representation for PHO84 has a strong activity for matrix 0, which is similar to the Pho4 consensus site. Each yellow bar represents the signal for each motif and brighter yellow indicates stronger signal, black indicates 0.

To detect conservation of motifs without sequence alignments, we first use a motif-based encoder to convert each set of orthologous promoter regions into a fixed-length representation vector (or “embedding”), where each element of the vector (or “dimension”) corresponds to a single motif[31]. We followed recent work[35] showing that motif match cutoffs, match counting and motif interactions can be modelled explicitly using trainable scaling, pooling and attention layers, respectively (Figure 1b). Standard approaches to model motifs in bioinformatics (position weight matrices, or PWMs[36]) are based on probabilistic models, so we ensured that the model learns valid PWMs by constraining the probabilities at each position to add up to 1 (see Methods). The resulting biologically interpretable neural network[35] embeds (unaligned) individual sequences in a space of 256 learned motifs.

To infer the parameters of the interpretable encoder (PWMs, scaling, interactions), we used reverse homology ([38], see Methods). This is a contrastive learning approach that uses orthologous sequences alone to develop a representation that maximizes mutual information between sequences of shared ancestry[37], which can be considered an alignment of orthologs in the learned representation space (instead of aligning in sequence space, standard in comparative genomics[8]). In practice, in order to learn similar representations of orthologs and different representations of unrelated sequences, reverse homology entails training a model to classify orthologs vs. non-homologous sequences by comparing the individual sequence representations to an estimate of the phylogenetic average representation of their orthologs (see Methods). In the case of our motif-based encoder, this encourages learning of motifs that can be identified in individual sequences and match the conserved motifs in the orthologs. Once the parameters of the encoder have been learned, we measure evolutionary conservation in the motif representation space simply by averaging each of the motifs’ scores over all the orthologs (Figure 1b). We refer to this as phylogenetic averaging[31], and it allows us to detect conservation of motifs in the representation space, rather than in sequence alignments, greatly increase the evolutionary diversity of the sequences we can include. Thus, using reverse homology (RH) we train an interpretable encoder (IE) and obtain a representation for a set of orthologous upstream regions by phylogenetic averaging (PA). We abbreviate this as RHIEPA and show an example of the representation of *PHO84* in Figure 1c.

To first test the power of motif-based embedding learned through reverse homology (RHIEPA) on a well-studied regulatory network, we obtained orthologous upstream sequences for *S. cerevisiae* promoters from 89 species from the y1000+ project[24] (see Methods) yielding nearly 380,000 non-coding sequences. This represents an order of magnitude more data than classical comparative motif-finding approaches[9], but an order of magnitude less than used to train large-language-models[41] (See Discussion).

We found that many of the PWMs learned by the model agree with known motifs for TFs from the JASPAR fungi database [42] (Figure 2a). These include both long, information-rich motifs that have high signal-to-noise (Figure 2a, matrix 81/Sfp1, matrix 66/Mcm1), but also short motifs with little information content (Figure 2a, matrix 29/Crz1). Like known transcription factor binding motifs, the learned PWMs showed overlapping sequence preferences (Figure 2a, matrix 121/Swi4, matrix 83/Mbp1::Swi6). Overall, 63 of the 256 learned PWMs match known motifs when we compare to two current databases ([42,43], Figure 2b) using the MEME Suite’s TomTom motif database search tool ([44], E-value < 4e-03, see Methods). 63 known motifs is nearly half of the number that match (E-value < 4e-03) when we search the same databases using a high-confidence, expert curated set of PWMs for yeast [45] (based on 100s of *in vitro* and *in vivo* experiments related to transcription factor specificity) (Figure 2b). This indicates that the motifs learned from sequences alone contain a reasonable fraction of the knowledge about the cis-regulatory vocabulary that has been learned from decades of direct experiments.

**Figure 2.**
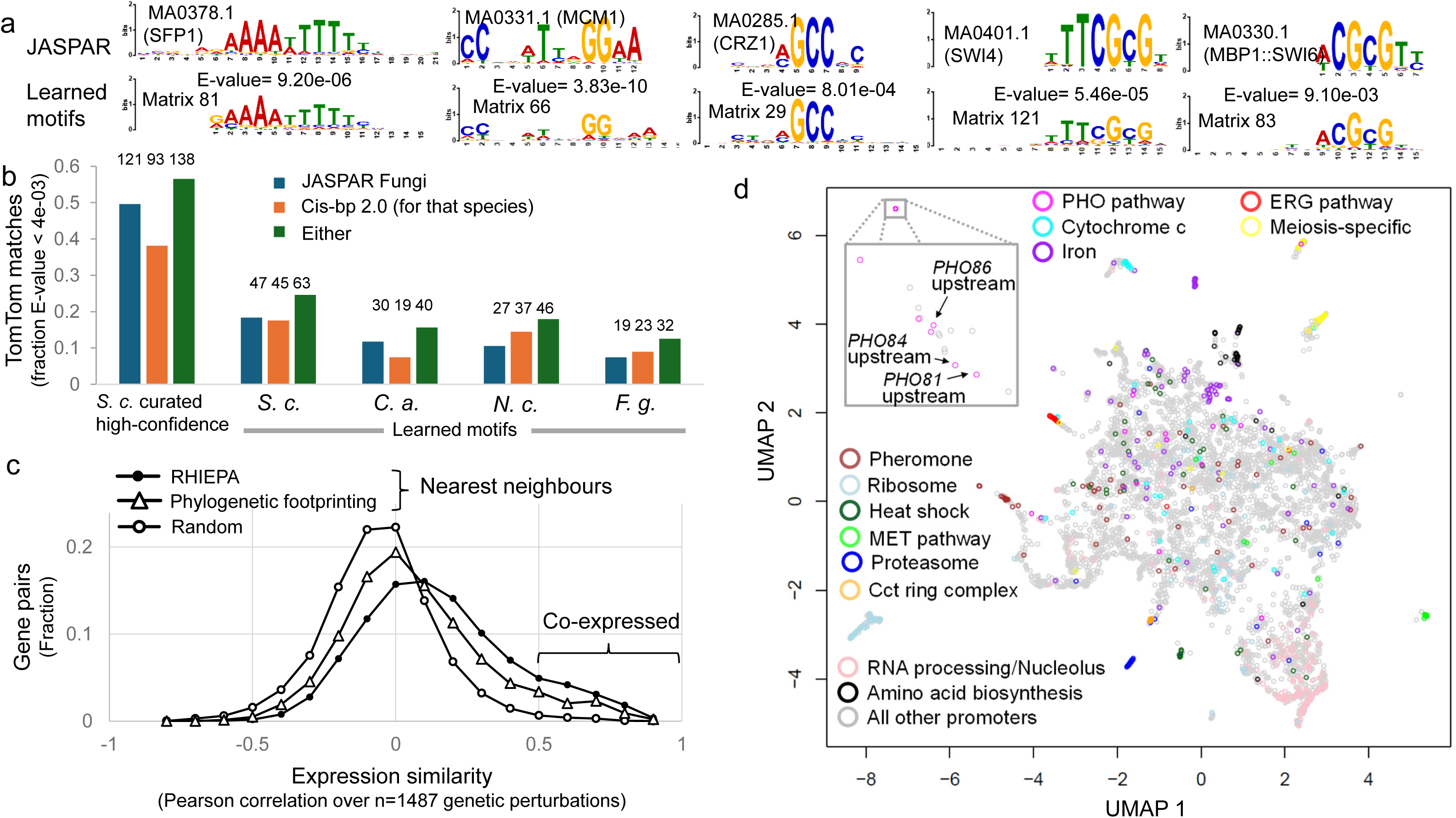
Unsupervised learning of the cis-regulatory code from sequence alone. a) Examples of learned yeast motifs (Matrix x) with matches to motifs the JASPAR database (MAx.x). E-values (from TomTom) indicate the probability of observing a motif in the database as similar by chance b) summary of learned motif matches against JASPAR and Cis-bp databases for 4 clades considered here (S. c.: *S. cerevisiae*, C.a: C. albicans, N.c: *N. crassa* F.g.: *F. graminearum*. High-confidence motifs from the Yetfasco database are included as a positive control for the database search. Numbers above the bars indicate the number of matches. c) gene expression similarity of nearest neighbours the <u>r</u>everse <u>h</u>omology, interpretable <u>e</u>ncoder, <u>p</u>hylogenetic <u>a</u>verage (RHIEPA) promoter sequence representations (filled symbols) compared systematic phylogenetic footprinting (triangles) and random neighbours (unfilled circles). Expression similarity is measured using Pearson correlation over 1487 genetic perturbations. When a pair of genes shows correlation greater than 0.5, we consider them to be co-expressed. d) biological organization emerges in a 2D representation of the 256 dimensional RHIEPA space for *S. cerevisiae*. Each symbol represents a set or orthologous upstream regions. Colours indicate matches to the *S. cerevisiae* gene name or description. Inset is a zoom in showing that the PHO84 upstream region is found among upstream regions from other genes in the PHO pathway.

To confirm that averaging the motif scores from the interpretable encoder over all the homologs (so-called phylogenetic averaging[31]) predicts TF targets, we selected 15 transcriptional activators for which a motif in the JASPAR fungi database was similar to the RHIEPA learned motif (TomTom E-value<0.05) and for which there was TF overexpression time-course data [46]. For each of these, we obtained the promoters bound in vivo according to the Yeastract database [47] and considered the genes with median log expression change >1 in the TF overexpression time course to be responsive to the activator over-expression (Supplementary Figure 1a). To ensure a fair comparison, we selected the promoters with the highest RHEIPA signals for each TF (when multiple learned motifs matched the TF, we used the median score; see Methods), choosing the same number of promoters as were bound in vivo. We found that the RHIEPA predicted targets (which are based only on genome sequences) were significantly more likely to respond to overexpression of the corresponding TF than the in vivo binding targets (15% vs. 10% P=0.037, paired t-test, n=15). These results indicate that conservation of learned motifs can be as good as or better than in vivo binding experiments at identifying promoters that respond to specific TFs.

An important potential problem in using genome sequences to predict regulatory networks is that TFs may bind to regions of the genome without matches to their motifs[48]. At least in this data, we find binding events without RHIEPA motif signals are rarely associated with gene expression responses. Over the 15 activators considered here, on average 92.6% (s.e.=1.4%) of the in vivo bound promoters that responded to TF overexpression were among the top RHIEPA predicted TF targets.

We wondered whether the low fraction of responsive genes among the in vivo bound targets (low precision or positive predictive power) was due to lack of responses (or responses below our cutoff) of *bona fide* target genes in the TF overexpression experiments. To test this, we looked at the top 50 promoters as ranked in the RHIEPA representation (Supplementary Figure 1a, green bars). Inconsistent with lack of responses in expression experiments (or too stringent a cutoff for determination of responses), we found a much higher positive predictive power: on average 40% (s.e.=6%) of the top 50 promoters responded, more than twice the fraction at the number of targets found in Yeastract (P=0.002, paired t-test, n=15). This suggests that the poor predictive power when considering the large numbers of targets found in Yeastract is likely due to the previously reported observation that that TFs bind many sites in vivo that are not associated with gene regulation [48,49]. Indeed, for many of the transcription factors, the number of in vivo bound targets in yeastract is in the 100s or 1000s (Supplementary Figure 1a), much greater than the number of genes that are involved in the associated biological pathways.

### Location in the learned motif space is associated with gene expression patterns and gene function

As gene expression control is appreciated to result from combinations of multiple transcription factors, we next sought to assess the biological information in the combination of all motifs within the high-dimensional motif-based phylogenetic average representations of promoter regions (RHIEPA, Figure 1c). To do so, we first compared the promoter representations to gene expression patterns over more than 1400 yeast gene deletion experiments[50]. Since functionally similar promoters are expected to drive similar expression patterns, we identified nearest neighbours (using cosine distance, see Methods) for each promoter in the RHIEPA representation space. We quantitatively evaluated the similarity of expression patterns driven by neighbouring promoters in two ways. First, we tested how well we could predict gene expression in each of 1487 experiments by using the expression of the gene downstream of the nearest neighbour promoter as the prediction (1-NN regression, see Methods). Second, we used the neighbouring pairs of promoters to identify co-expressed pairs of genes (defined as gene pairs with correlation >0.5 over the 1487 experiments). These represent measures of prediction of between-gene and between-condition variation, respectively[51], based on the promoter sequence alone. We emphasize we did not fit any parameters using the gene expression data or include any prior knowledge to make predictions. This is in contrast to current regulatory network inference approaches for well-studied cells that leverage prior knowledge about transcription factors, their motifs, as well as large-scale data for gene expression, chromatin and/or transcription factor binding [52–54]. We compared the predictions of gene expression and co-expression to predictions using nearest neighbours obtained from a systematic phylogenetic footprinting approach[39] applied to our data (conservation tests for 41,600 k-mers, see Methods) or from the regulatory network inferred by NetProphet3[54] (an approach to regulatory network inference that combines multiple types of experimental data, see Methods). Nearest neighbours in the RHIEPA representation space (Figure 2c, filled symbols) are more likely to show similar gene expression patterns than nearest neighbours in the systematic phylogenetic footprinting space (Figure 2c, triangles) and far more likely than expected for random neighbours (Figure 2c, unfilled circles).

Averaging over all the gene expression experiments (Table 1), RHIEPA representations of promoters predict gene expression with R=0.221 (s.e. 0.003), greater than R=0.122 (s.e. 0.001) for NetProphet3, and far greater than R=0.056 obtained using the top BLAST hits (see Methods). Similarly, over all genes, the RHIEPA nearest neighbour is co-expressed for 768 genes (14.3% or 0.14 precision, n=5353 genes total, Table1), more than the 236 genes (4.4% or 0.04 precision, n=5364 genes total, Table 1) for NetProphet, and the 213 genes (4.0%, n=5303) observed for best BLAST hits. Because the classification of co-expressed gene pairs is outside of the training data of the sequence-based RHIEPA, this is a measure of “zero-shot” generalization.

**Table 1.** comparison of S. cerevisiae promoter representations to gene expression data. Representation dimensionality indicates the length of the numerical description for each promoter. For Netprohet3 and Yeastract, these are binary (yes or no) whether a transcription factor (TF) binds to the promoter of that gene. Best matches in S. cerevisiae and phylogenetic average motif scores use normalized matches to the motif for each transcription factor from the yetfasco database. Phylogenetic footprinting uses a log P-value from the binomial distribution for each of the 41,600 spaced k-mers (or dyads) that has been corrected and thresholded, so non-significant values are 0. RHIEPA uses the signals from the 256 learned motifs. Methods are grouped by the required input data: direct experimental measurements of transcription factor-DNA interactions vs. transcription factor binding motifs obtained from in vivo or invitro DNA binding experiments (but no in vivo targets needed) vs. genome sequences from multiple species (no information about transcription factors or their motifs is needed). Left two columns represent nearest neighbour regression (nn) to predict relative gene expression levels. Right panel represents precision (positive predictive power) to identify a co-expressed gene as the closest neighbour (Top 1) or among the 50 nearest neighbours (Top 50). Bold text indicates the best performing method. Error bars represent 3 times standard error of the mean over n=1487 correlation coefficients (one for each experiment) or using the normal approximation to the binomial for the precision (n>5300 genes). Three standard errors were used as a correction for multiple testing because of the large number of comparisons presented.

| Approach<br>(representation<br>dimensionality) | Expression experiment nearest-neighbour<br>regression (average Pearson correlation<br>over n=1487 deletions strains, +/- 3 s.e.) |  |  |  | Gene co-expression prediction (Zero<br>shot precision over n>5300 genes, +/-<br>3 s.e.) |  |  |  |
| --- | --- | --- | --- | --- | --- | --- | --- | --- |
|  | 1-nn |  | 50-nn |  | Top-1 |  | Top-50 |  |
| Use direct experimental data for TF-promoter interactions |  |  |  |  |  |  |  |  |
| Netprophet3<br>(185 TFs) | 0.122 | +/-0.004 | 0.177 | +/-0.005 | 0.044 | +/-0.008 | 0.310 | +/-0.019 |
| Yeastract (182<br>TFs) | 0.096 | +/-0.003 | 0.167 | +/-0.005 | 0.044 | +/-0.008 | 0.336 | +/-0.019 |
| Use known motifs, but no direct measurement of interactions |  |  |  |  |  |  |  |  |
| Best matches in<br>*S. cerevisiae*<br>(244 motifs) | 0.073 | +/-0.003 | 0.125 | +/-0.006 | 0.053 | +/-0.009 | 0.306 | +/-0.018 |
| Phylogenetic<br>Average Motif<br>Scores (244<br>motifs) | 0.174 | +/-0.007 | 0.277 | +/-0.008 | 0.118 | +/-0.013 | 0.417 | +/-0.020 |
| Use genome sequences only |  |  |  |  |  |  |  |  |
| Phylogenetic<br>footprinting<br>(41,600 k-mers) | 0.138 | +/-0.006 | 0.245 | +/-0.008 | 0.088 | +/-0.011 | 0.357 | +/-0.020 |
| RHIEPA (256<br>learned motifs) | **0.221** | +/-0.008 | **0.344** | +/-0.009 | **0.143** | +/-0.014 | **0.453** | +/-0.020 |

To further understand the improved association with gene expression of the RHIEPA representation, we tested a representation based on the phylogenetic average of motif scores [31] of expert curated PWMs[45] and found that it also yields better predictions of expression (1-NN regression R=0.174 +/- 0.002) and co-expressed gene classification (Zero shot precision 0.12) than systematic phylogenetic footprinting, NetProphet3 neighbours and experimental binding data (Table 1), nearly as high as the RHIEPA representation (1NN regression R=0.221, Zero shot precision 0.14, Table 1). However, we find that matches to the expert curated PWMs in *S. cerevisiae* alone (without phylogenetic averaging) have little power (Table 1), consistent with the idea that matches to motifs in single genomes are mostly non-functional noise (the so-called futility theorem[4]). Since single nearest neighbours explores only highly local relationships in the high-dimensional representation space, we repeated these experiments for 50 nearest neighbours and found similar trends (although with overall greater predictive power, Table 1). This indicates that the similarity in the motif-based representation space extends beyond pairs of promoters, as expected if the representations are grouping together co-regulated pathways. Taken together these results suggest that much of the power to predict gene expression (relative to TF binding data and methods based upon it) is due to the phylogenetic averaging, which we believe increases the signal-to-noise ratio[31].

We next visualized the RHIEPA representation in 2 dimensions using UMAP (see Methods), where each point corresponds to the 256-dimensional vector for a promoter (and its orthologs, Figure 1c). We coloured the points representing promoter regions matching English text in the associated gene names or descriptions (see Methods) and found that promoter regions flanking genes with the same matches to their names and descriptions were nearby each other (Figure 2d). For comparison, we computed a similar UMAP for the representation from phylogenetic footprinting (Supplementary Figure 1b). We emphasize that only genomic sequence information is used in training the model or making the UMAP, and all the orthologs for each promoter have been averaged into a single point (proximity of promoters is not due to sequence similarity between orthologs), and the colouring is not based on clustering. The proximity of promoters with the same text in their gene names and descriptions is further evidence that the motif-based representation contains functional representations of the promoters.

Taken together these results show that training a motif-based representation on genome sequences alone identifies known motifs, predicts gene expression patterns and groups promoters according to their biological functions.

### Reading the cis-regulatory code by visualizing the motif-based representation

Given the associations of the RHIEPA representation with gene function and gene expression, we next sought to explore whether we could understand how the presence of motifs determines promoter function (the cis-regulatory code). We clustered and visualized[55] the *S. cerevisiae* promoter regions based on their RHIEPA representations (Figure 2a). For exploration of the clustered heatmaps, we named learned motifs using TomTom matches to the JASPAR fungi database (E-value < 0.05, See Methods). We then identified patterns of motifs that were strongly associated with GO biological functions (using Gprofiler[56] enrichment analysis and corrected P-values, Supplementary File 1). For example, consistent with known regulation [57], a cluster of 265 promoters with strong signals for Dot6, Stb3, Sfp1 and Abf1 is strongly enriched for genes annotated as ribosome biogenesis (162 of 263 total annotated genes, 9.6x enrichment, P= 5.45E-133, Figure 3a) and is distinct from a cluster of 120 promoters that does not show signals for Stb3, but does show signals for the other factors, and is associated with genes annotated as translation machinery (61/119 total annotated, 6.8X enrichment, P=4.09E-35, Figure 3a). This cluster includes promoter regions for t-RNA synthetases, which additionally show a signal for a motif matching Gcn4 (known to regulate amino acid biosynthesis[58]) consistent with a recent report of activation of Gcn4 when tRNA synthetase activity is low[59] and more generally, a connection between regulation of amino acid biosynthesis and translation. Also consistent with known regulation[57], a cluster of 124 promoters associated with ribosomal proteins (82/124, 24x enrichment, P=3.10E-104, Figure 3a) are distinguished by signals for a learned motif matching Rap1. Finally, a cluster of promoters associated with genes annotated as mitochondrial translation (36/260, 6.7x enrichment, P=7.56E-18, Figure 3a) also shows signals for Sfp1 and Abf1, but also Nsi1 (which is similar to Reb1). Thus, the combinatorial control of translation and associated cellular machinery can be read from this visualization of the motif-based promoter representation (see Discussion).

**Figure 3.**
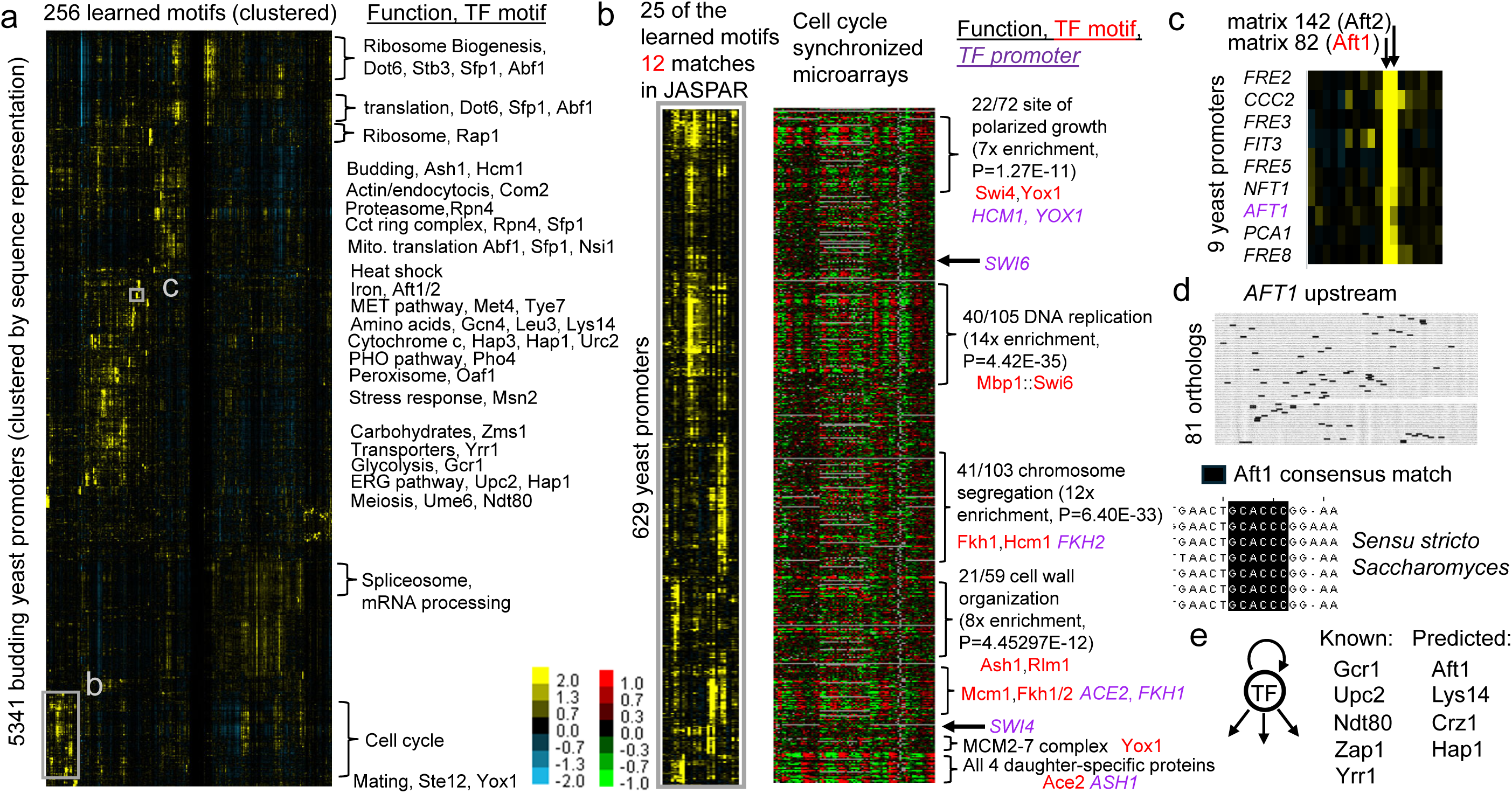
Reading the cis-regulatory code in yeast. Reading the cis-regulatory code in yeast. a) yeast promoters can be organized by sequence alone one they are transformed into the motif-based RHIEPA representation. Rows represent orthologous sets of upstream regions and columns represent learned motifs. Yellow colour indicates stronger signal for that motif in that promoter. Biological functions and known transcription factor (TF) motifs (best match of the learned motif JASPAR, TomTom E-value<0.05) are indicated to the right of clusters. Grey boxes in the heatmap indicate clusters in subsequent panels. b) the upstream regions of genes encoding cell cycle transcription factors (purple) are in a cluster where the learned motifs match their known motifs (red). Gene expression timecourses from yeast cells synchronized by alpha factor arrest, cdc15 or cdc28 mutants or elutriation is ordered according to the clustering of motif-based promoter representations and red indicates higher relative expression, while green indicates lower relative expression. GO enrichments for smaller clusters are indicated to the right of the heatmaps. c) a small section of the clustered promoter heatmap shows the promoter for AFT1 among known iron-responsive genes. d) Aft1 is predicted to be autoregulated because the upstream regions of AFT1 and its orthologs contain matches to the Aft1 consensus site, and one of these sites is conserved among closely related yeasts. e) transcription factor (TF) autoregulation is predicted based on the promoter representation cluster analysis in at least 9 cases, 5 of which are known (see text for details)

To further explore whether the signals in the RHIEPA representation show evidence of combinatorial regulation, we next examined a large cluster of *S. cerevisiae* promoters showing signals for many of the known cell cycle regulatory transcription factors and enriched for promoter regions upstream of cell cycle genes (262/629 promoters, 3.2x enrichment, P=1.93E-79). Comparing this cluster of promoter regions to gene expression measurements for the associated genes (Figure 3b) revealed that the groups of promoters with signals for different patterns of TFs correspond to groups of genes with coordinated gene expression patterns in classic cell cycle synchronization experiments[60] (Figure 3b, full comparison in Supplementary Figure 2a) and correlated cell heterogeneity in sc-RNA-seq[61] (Supplementary Figure 2b).

These groups of genes showed different functional enrichment (Figure 3b) consistent with regulation by different combinations of cell cycle TFs that control different stages of the cell cycle. For example, a cluster of promoters with signals for learned motifs matching Mbp1::Swi6 shows enrichment for genes involved in DNA replication (Figure 3b), while a cluster with signals for motifs matching Hcm1 and Fkh1 shows enrichment for genes involved in chromosome segregation (Figure 3b). These observations are consistent with the distinct cell cycle functions of these TFs[62,63]. To get a quantitative sense for the power of the clustering approach to identify genes with complex regulation, we compared this cell cycle cluster to a curated set of cell cycle and non-cell cycle genes used to train machine learning classifiers [64]. We found that our cluster gives a true positive rate of 48% and a false positive rate of 4%, comparable to the predictive performance of machine learning classifiers trained on this dataset using selected *in vivo* binding experiments and matches to known motifs (Supplementary Figure 2c). Thus, at least in this well-studied case, we find that the procedure of identifying clusters with strong signals in the RHIEPA representation (which is made from genome sequences alone) identifies genes regulated by multiple TFs as well as machine learning classifiers trained using curated labels and experimental data to do this task.

We also observed that TF-TF regulatory connections could be inferred while analyzing this cluster of promoters associated with the cell cycle: of the 12 learned motifs that matched JASPAR (Figure 3b, red text), the promoters of genes encoding 8 (66%) of the corresponding TFs (Figure 3b, purple text) were found in the cluster. Since less than 15% of the genes in the genome are in the cluster, this represents a strong statistical enrichment (P=1.23E-05, Fisher’s exact test) and is consistent with the known regulation of the cell cycle TFs by each other[65].

Further supporting the idea that TF regulation is visible in the sequence representation, we found we could infer examples of autoregulation in the *S. cerevisiae* promoter map. For example, the promoter region of the gene encoding the iron-responsive transcription factor Aft1[66] was found among promoters upstream of genes encoding transition metal transporters (31/47, 60x enrichment, P=1.19E-49) in a cluster with strong signals for learned motifs that matched the Aft1 and Aft2 motifs in JASPAR (Figure 3c shows a subset of this cluster).

Although to our knowledge auto-regulation of *AFT1* has not been reported, the promoters of *AFT1* orthologs contain matches to the Aft1 consensus (Figure 3d), consistent with binding of Aft1 to its own promoter. This prediction is also consistent with a peak of Aft1 binding observed at the AFT1 promoter (Supplementary Figure 2d) under standard growth conditions [67]. More generally, we found at least 9 (Figure 3e) examples of TF promoters showing strong signals for the learned motifs that matched their own motifs in JASPAR and were clustered among their known target promoters in the motif-based sequence representation. Of these, at least 5 are known (Gcr1[68], Upc2[69], Ndt80[70], Zap1[71] and Yrr1[72]), while 4 (Aft1, Lys14, Crz1 and Hap1) appear to be new predictions of TF autoregulation.

Taken together, the cluster analysis of the phylogenetic averages of learned motif representations for the well-characterized regulatory network of budding yeast supports the idea that we can infer regulatory networks from genome sequence alone.

### Prediction of a meiosis/parasex pathway in C. albicans

Because the regulatory inference based on genome-sequences is unbiased by experimental conditions or lab constraints, we next tried to infer a regulatory network for *C. albicans*, a distantly related pathogenic fungus with many shared genomic features to *S. cerevisiae*. We obtained orthologous promoter sequences for *C. albicans* from 69 species from the Y1000+ project (see Methods) again yielding more than 300,000 non-coding sequences. We inferred the parameters of interpretable encoder using reverse homology with no changes to hyperparameters and created representations for the promoters by phylogenetic averaging (RHIEPA). Consistent with the relative paucity of experimental measurements in *C. albicans*, many fewer of the learned motifs matched known motifs in the databases relative to *S. cerevisiae* (Figure 1b). Nevertheless, we observed many of the same patterns in the representation space, where clusters of promoters showed strong functional enrichments (Figure 4a, Supplementary File 1).

**Figure 4.**
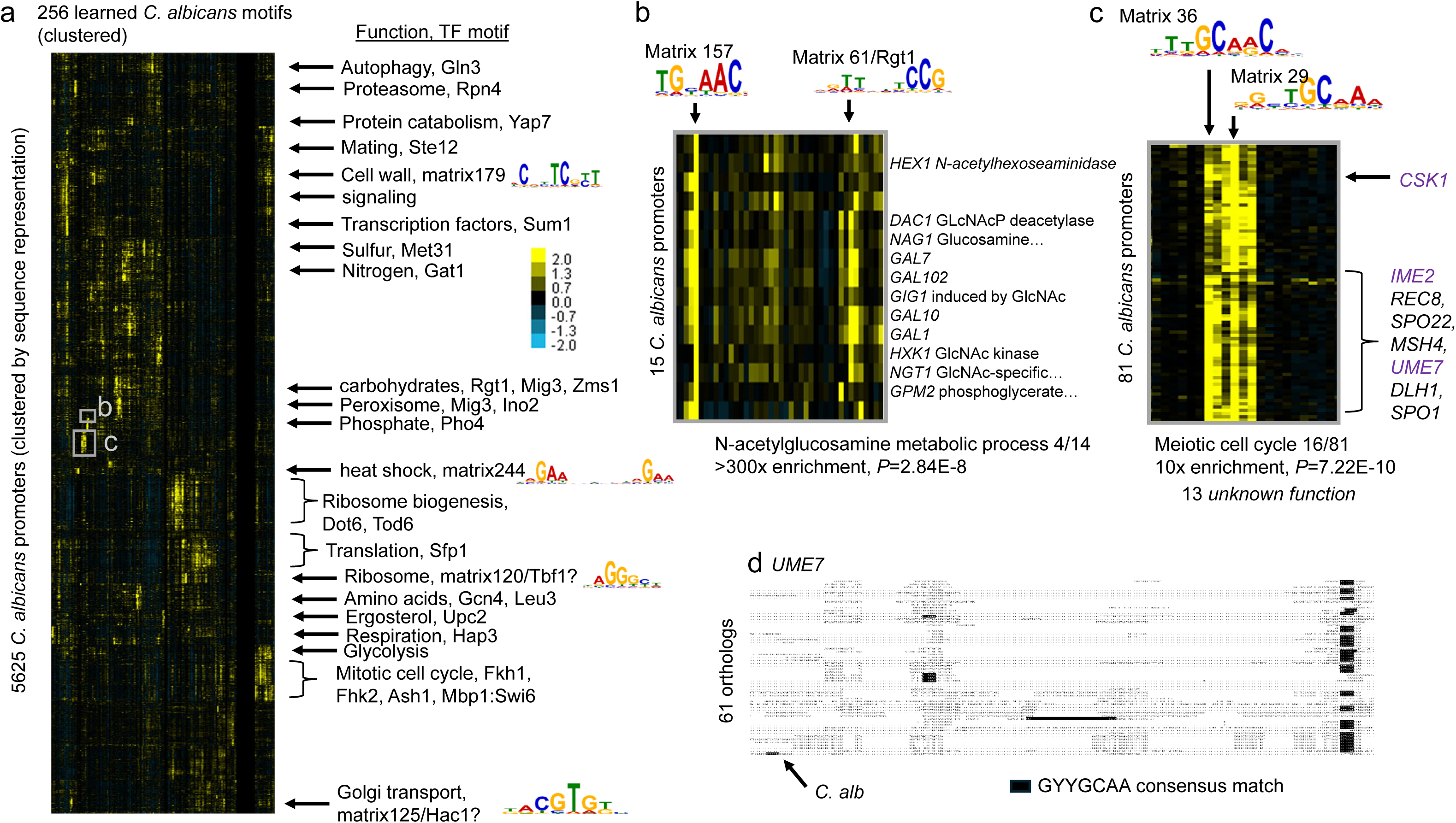
A regulatory map for C. albicans predicts a meiosis/parasex pathway. a) clustered motif-based RHIEPA representations of *C. albicans* promoter regions presented as in Figure 3a. TF motifs with ‘?’ did not have a significant match in JASPAR, but are named based on similarity of the logo (displayed) to known motifs. b) a cluster of promoters associated with the GAL genes (*GAL1,GAL10,GAL7,GAL102*) as well as several genes related to GlcNac (*DAC1, NAG1, GIG1, HXK1, NGT1*). Genes without names are left as blank rows. Two learned matrices with strong signals in these promoters are indicated above. c) cluster of upstream regions statistically associated with meiosis annotations. Upstream regions of homologs of meiosis genes are indicated in black, while upstream regions of homologs of meiosis regulators are indicated in purple. Two learned matrices with no matches to JASPAR (Matrix 36 and Matrix 29) show strong signals in this cluster. d) part of the alignment of the 61 orthologous *UME7* upstream regions. *C. albicans* sequence is indicated at the bottom. Black boxes indicate matches to the consensus sequence of the learned motifs shown in c).

We also noted some known evolutionary differences in regulation between *C. albicans* and *S. cerevisiae*, such as different motif signals in a cluster of promoter regions of glycolysis genes[73] (8/14, >180x enrichment, P=2.0E-15) and absence of Rap1 motifs in promoter regions associated with the ribosome genes [74] (63/76, 34x enrichment, P= 9.33E-92).

Although it has no significant match in JASPAR fungi or cis-bp2.0, a learned motif (matrix 120) associated with the promoters of the ribosome cluster matches the reported consensus for Tbf1[75] (Figure 4a), which is expected to regulate ribosome gene expression in *C. albicans* (along with Sfp1 and Stb3) [74]. In a more complicated example, we find a cluster of promoters including both the well-known *GAL* genes, and genes involved in GlcNAc signaling (Figure 4b) that were recently found to be co-regulated by Rep1[76]. The motif identified (matrix 157) resembles the half-site previously identified in *C. albicans GAL* promoters[77], so we propose that it represents the binding site for Rep1. We also noted the presence of signals for a motif (matrix 61) with a strong similarity to Rgt1 in JASPAR fungi (TomTom E=7.60e-07) in these promoters. Based on this, we predict additional regulation by Rgt1 of the GAL and GlcNAc genes in *C. albicans*.

Unexpectedly, we identified a group of promoter regions strongly associated with genes annotated to have a function in the meiotic cell cycle (Figure 4c). Meiosis has never been observed in *C. albicans*, although *C. albicans* is known to have several of the homologs of *S. cerevisiae* meiosis genes and undergoes a recombination process known as parasex[78]. In this cluster, we also found upstream regions of homologs of yeast meiosis regulators (*UME7*, *CSK1* and *IME2*). The motifs with strong signals in this cluster (Figure 4c) did not match any known motifs in JASPAR, but we find clear conservation in promoter sequences (Figure 4d). We suggest that they represent the regulators of parasex in *C. albicans*, and the (at least) 13 genes of unknown function in this cluster are candidates for genes involved in meiosis/parasex.

### Inferring regulatory networks for multicellular fungi

Encouraged by the analysis of unicellular fungi, we next attempted to apply this approach to more complex organisms. *Neurospora crassa* is a genetic model system for filamentous fungi. Its genome is 40MB (compared with 12MB for budding yeast) and contains approximately 10,000 genes (compared with approximately 5,000-6,000 for budding yeast) nearly half of which are annotated as encoding unknown proteins[79]. We obtained orthologs from the 52 closest genomes (see Methods), again yielding more than 300,000 non-coding sequences and with no changes to hyperparameters created a motif-based RHIEPA representation. The motifs learned matched more known motifs in the Neurospora-specific cis-bp database than JASPAR fungi (37 vs. 27) consistent with many of these specificities being predicted from *in vitro* TF binding experiments[43].

As with yeasts, we found that clusters of promoter regions with similar signals in the RHIEPA representation showed strong association with essential cellular functions (Figure 5a). For example, we observed a signal for a matrix that matches JASPAR fungi Gat1 in a cluster of promoters associated with nitrogen transport (30/189, 3.2x enrichment, P= 7.99E-06), consistent with previously observed conserved regulation[80]. As in yeasts, we find separate clusters of upstream regions for ribosome biogenesis genes (109/233, 18x enrichment, P=8.66E-113), translation factors (19/68, 39x enrichment, P=7.86E-24) and ribosome components (53/81, 38x enrichment, P=2.98E-74). The ribosome components include a signal for a similar Tbf1-like motif as found in *C. albicans*, while Ribosome biogenesis promoters include signals for Sfp1 (as in *S. cerevsisiae*) but also include signals for a GATA factor motif that matches Gat1, suggesting different mechanisms may regulate growth in Neurospora.

**Figure 5.**
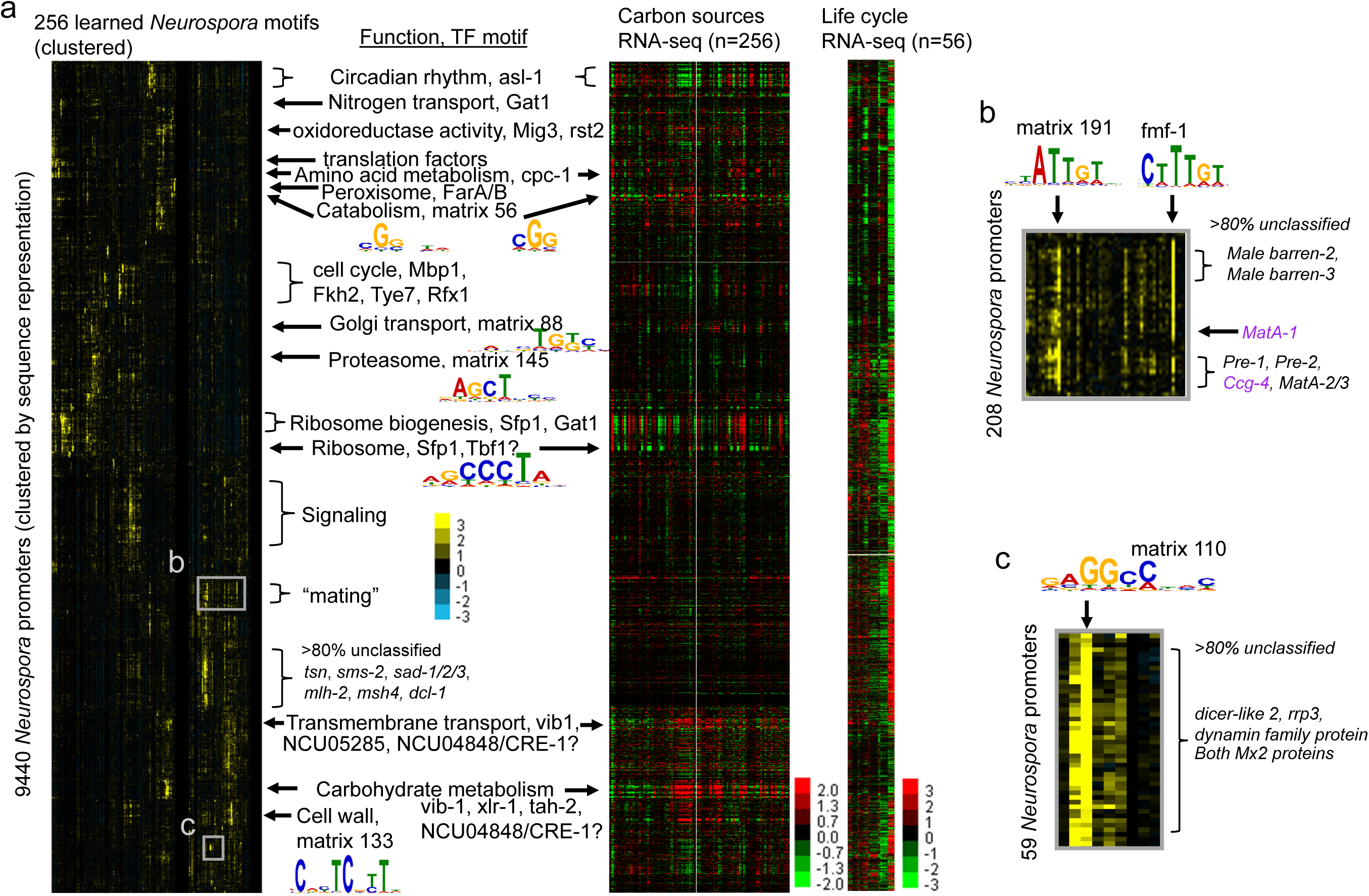
Reading the cis-regulatory code in N. crassa. a) heatmap of motif-based RHIEPA promoter region representation and comparison with gene expression is shown as in Figure 3a. Matrices with no matches are indicated as matrix x. question marks indicate motifs that are similar to known motifs, but not statistically significant in database search. Predicted b) mating pathway and c) virus response pathway displayed as in Figure 4b. Ccg-4 is the only known target of MatA-1 (purple).

We find a cluster of ∼500 promoters strongly associated with the cell cycle (163/505, 6.1x enrichment, P= 1.16-86) that share some motifs with yeast (Mbp1[80] and Fkh2) alongside signals for different motifs (Tye7 and Rfx1), suggesting different regulation. In contrast to the well-studied budding yeast cell cycle, 40.6% of the genes in the cell cycle cluster in *Neurospora* are annotated as encoding “hypothetical protein” and little is known about the transcription factors that control the *Neurospora* cell cycle. Thus, our analysis provides hypotheses about the identity of cell cycle regulators in this species.

Other clear signals include a cluster of promoters strongly associated with peroxisome genes (21/96, 32x enrichment, P=9.57E-25) shows a strong signal for matrix 234 that has no matches in the databases, but matches the CCGAGG consensus of FarA/B, transcription factors that regulate the peroxisome genes in Aspergillus[81]. Furthermore, the promoters of proteasome subunits contain signals for unknown motifs (matrix 145, Figure 5a) consistent with previous reports that transcriptional regulatory of the proteasome has diverged in fungi[80]. Similarly, upstream regions associated with Golgi transport, cell-wall components and genes involved in catabolism also show unknown motifs (matrix 88, matrix 56, matrix 133, Figure 5a). These represent new hypotheses for basic regulatory mechanisms.

Beyond basic cellular mechanisms, we identified a group of 165 promoter regions with signals for a matrix that matched ASL-1, a regulator of osmotic stress and circadian rhythm gene expression [82]. The genes associated showed no functional enrichment (though 55% are annotated as hypothetical protein) but do include 3 of the 7 known ASL-1 targets[83] (cat-1, bli3, and ccg-1, 27x enrichment, P = 0.00012, Fisher’s exact test). To test for circadian rhythm gene expression, we counted genes with both significant RNA and RNAPII-S2P rhythms [84]. We found 16 of 67 (15x enrichment, P = 2.85e-15, Fisher’s exact test) associated with the promoter regions in this cluster, indicating that many of the promoters in this group may be uncharacterized direct targets of ASL-1.

To confirm that similar promoters in the RHIEPA representation space showed similar gene expression patterns, we next compared the sequence-based clustering of Neurospora promoters with large-scale gene expression measurements for growth on varying carbon sources[85] (Figure 5a). Indeed, groups of promoters with motif signals matching known carbohydrate regulators (vib-1, xlr-1 and NCU04848, whose motif is similar to CRE-1[85]) correspond to groups of genes with strong induction of gene expression and the genes associated with these promoters are also strongly enriched for genes involved in carbohydrate metabolism and transmembrane transport (10x and 7.4x enrichment, P=6.74E-51 and P=1.82E-28, respectively). We also noted strong expression changes for the ribosome and ribosomal biogenesis genes, perhaps consistent with differences in growth on the different carbon sources (Figure 5a). Interestingly, the cluster of promoters associated with rhythmic expression shows somewhat reciprocal gene expression changes (Figure 5a). Although it is tempting to speculate that variation in carbon source may influence circadian transcription in Neurospora, we cannot rule out simple time-of-day effects.

Taken together these analyses show that we can identify known and predict new regulation for known genes in Neurospora, thus inferring a regulatory network in a multi-cellular organism from genome sequences alone.

### Prediction of mating and RNA-virus response pathways in Neurospora

We next searched for possible new pathways among the clusters of upstream regions associated with large numbers of uncharacterized genes in Neurospora and found several compelling examples. We found a cluster of promoter regions associated with signals for a motif matching fmf-1 (Female and male fertility protein 1) and an unknown motif (matrix 191) (Figure 5b). Although >80% of the genes in this cluster have unknown function, it includes *matA-1*, *matA-2/3* (Mating-type proteins), both Male-barren genes and the pheromone receptor genes, *pre-1* and *pre-2* (Figure 5b). We therefore speculate that this represents the (largely uncharacterized) mating pathway in Neurospora (7/214 are annotated as response to pheromone, 15x enrichment, P=9.92E-05). Consistent with this, the *fmf-1* motif matches the CTTTG consensus proposed for MATA-1[86] and the group of genes includes *ccg-4*, the only confirmed target of MATA-1[83]. We also noted a group of ∼500 upstream regions with signals from unknown motifs associated with mostly uncharacterized genes (87% unclassified). This group includes several of the genes believed to be involved in recombination in Neurospora (e.g., *sad-1*[87], *sad-2*[88], *tsn*, *sms-2* [89], *mlh-2* and *msh4*), so we speculate that this represents the meiosis pathway. Consistent with this hypothesis, gene expression of both the putative mating and meiosis clusters is strongly upregulated during sexual development (last columns of life cycle RNA-seq[90], Figure 5a).

Perhaps the most interesting uncharacterized pathway is a cluster of promoter regions with signals for matrix 110 (Figure 5c). Although 87% of these genes are unclassified, they include dicer-like 2 and *rrp3* (an RNA-dependent RNA polymerase) that were both induced in response to viral infection[91]. We propose that this pathway represents an RNAi-based immune response. Like the putative parasex pathway in *C. albicans*, these results show how clusters in the motif-based promoter representation space lead to hypotheses about new pathways in difficult-to-study experimental conditions (see Discussion).

### A partial regulatory map for Fusarium

To explore how far these results can be extended, we attempted to infer regulation from a RHIEPA representation for a much less characterized organism, *Fusarium graminearum*, an important agricultural pathogen. We obtained orthologs for more than 12,000 genes from 68 species (see Methods), but noted that a large number of orthologous upstream sequences were very similar at the DNA level, possibly reflecting the large number of closely-related Fusarium strains that have been sequenced[92]. We therefore filtered the orthologs to remove very similar sequences (see Methods) and were left with nearly 280,000 upstream sequences.

Given the much sparser functional annotation of the *F. graminearum* genome, we found fewer clusters of promoter regions that were associated with known functions. Nevertheless, we observed functional enrichments similar to Neurospora for some aspects of basic cellular machinery (Figure 6a) such as Sfp1 signals in promoters of associated with the ribosome genes (40/80, 51x enrichment, P= 8.87E-59), FarA/B-like signals in promoters associated with the peroxisome (15/145 19x enrichment, P= 1.33E-19) and promoters with signals for several motifs, including one similar to CRE-1 associated with carbohydrate metabolism and transport (4.8x and 3.6x enriched, P= 8.11E-26 and 1.83E-32, respectively). As with Neurospora, we could predict unknown pathways: a large cluster of promoters, some of which show signals for a learned motif that matches fmf-1, includes the *F. graminearum* mating type proteins, as well as orthologs of several Neurospora genes involved in mating and meiosis (Figure 6a “mating/meiosis”).

**Figure 6.**
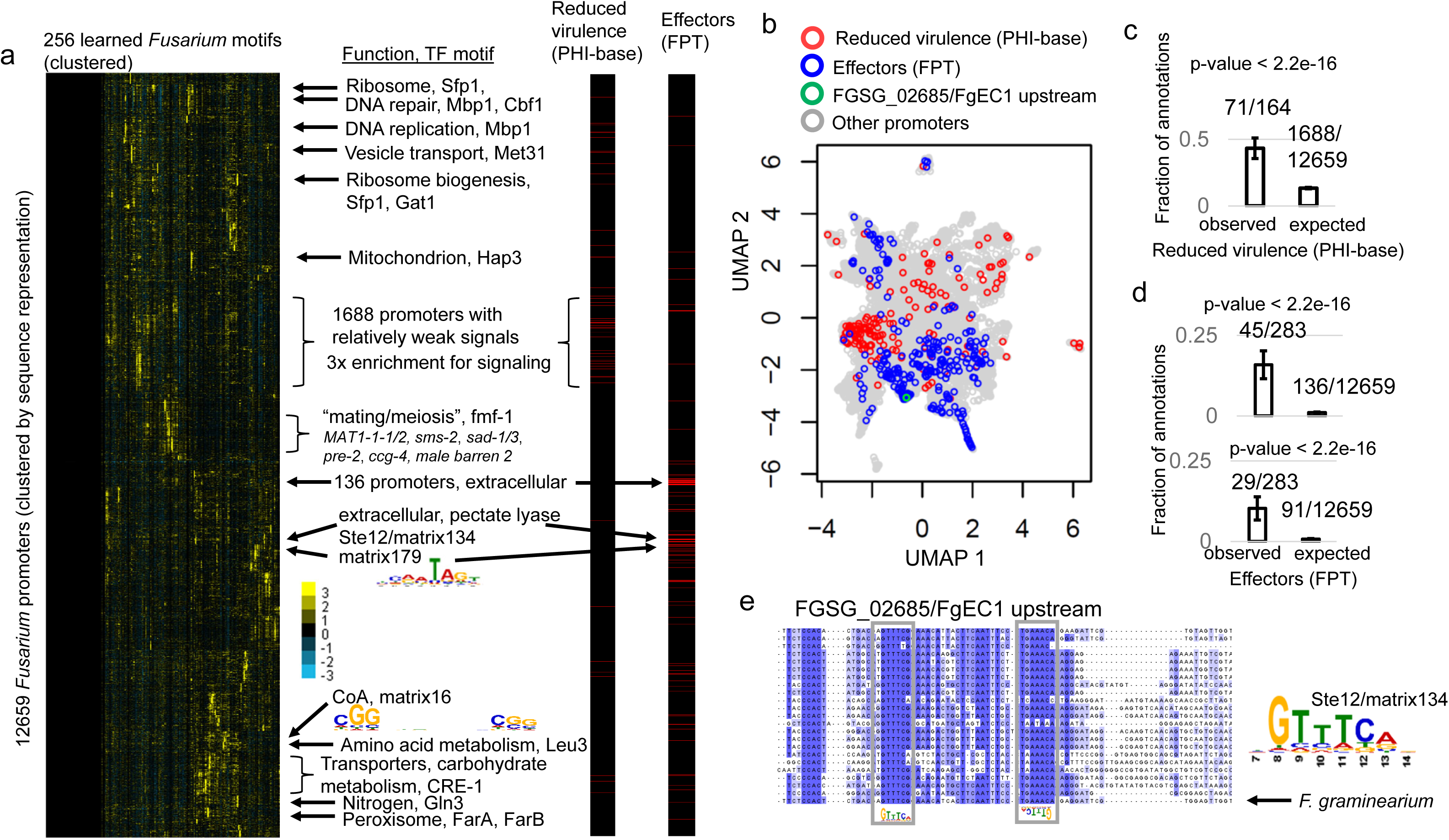
Association of promoter representation with virulence in Fusarium. a) Map of Fusarium promoter regions visualized as in Figure 3a shows many clusters with unknown motifs and no functional enrichments. A large group of upstream regions of genes enriched for signaling, but relatively weak motif signals is associated with reduced virulence annotations from PHI-base (red stripes in left vertical black bar). At least three clusters of promoters (indicated with arrows to red stripes in left vertical black bar) are associated with proteins encoding effectors (from Fusarium Protein Toolkit, FPT). b) promoters associated with genes annotated as reduced virulence (red symbols) are enriched in a different region of the high-dimensional motif-based representation than promoters associated with effectors (blue symbols). RHIEPA representation was reduced to 2D using UMAP, and promoters with neither annotation are indicated in grey. A green symbol indicates the location of the FgEC1 promoter. c) genes associated with promoters in the large cluster enriched for signaling are also associated with reduced virulence annotations in PHI-base. d) examples of clusters of promoters associated with effector proteins. Top panel shows a cluster of 136 promoters with no clear motif signals, while the bottom panel shows a cluster of 91 promoters with strong signals for matrix134 which matches Ste12 in a TomTom search. In c and d, error bars represent 2 times the standard error of the fraction (using the normal approximation to the binomial). Numbers above the bars represent the sample sizes used to compute the fraction. P-value is based on a Fisher’s exact test. e) portion of the alignment of FgEC1 ortholog promoters showing evolutionary conservation (blue highlight) of matches to a learned motif (grey boxes, Matrix134) that is similar to Ste12.

We next tested whether the RHIEPA promoter representation contained information about the pathways controlling virulence and pathogenesis in *F. graminearum*. To do so, we compared the motif-based clusters of promoters to annotations of “reduced virulence” phenotypes [93] and genes encoding predicted effector proteins[94] (Figure 6a, see Methods). The promoters for these two groups of genes showed little overlap (Figure 6b) indicating that they are associated with different regulatory mechanisms. We found that reduced virulence promoters were strongly enriched in a region of the RHIEPA representation space distinct from the promoters associated with basic cellular processes with relatively weak signals for learned motifs (Figure 6a), but a 3.8-fold enrichment for genes annotated as signaling (78/1650 of the 1688 promoters where we could map genes, P=4.62E-25). We found that the genes associated with promoters included 71 (43%) of the genes annotated as reduced virulence. Since this group represents 13% of the upstream regions in the genome, this is a highly significant enrichment (Figure 6c, Fisher’s exact test, P < 2.2e-16). Although it is tempting to speculate that promoter regions can predict virulence, given the enrichment of annotated signaling proteins in this group, we cannot rule out that the enrichment is due to bias in researchers to study the effects of mutations in signaling proteins on virulence.

We noted several small clusters that showed associations with effectors, the strongest of which was a 14.8x enrichment for a cluster of 136 promoters, associated with extracellular proteins (10.6x enrichment, P=1.8e-05) but no strong motif signals (Figure 6d, top panel, Fisher’s exact test, P < 2.2e-16). Another cluster strongly associated with effector proteins (14.3x enrichment, Figure 6d, bottom panel, Fisher’s exact test, P < 2.2e-16) was associated with genes annotated with GO terms extracellular region (35.8x enrichment, P=1.36e-25) and pectate lyase (93.1x enrichment, P=1.31e-14). Pectin lyases and their precursors are among the known effectors[94] and are important for pathogenicity *F. graminearium*[95]. This cluster of promoters showed signals for a learned motif matching Ste12 (Figure 6e), consistent with recent reports associating Ste12 with pathogenicity in Fusarium[96]. One of the genes in this cluster, FgEC1 (FGSG_02685/ FGRAMPH1_01G06433), increases expression during infection [97], is secreted, and interacts with wheat proteins. The strong signals for the Ste12 motif in its promoter correspond to conserved binding motifs (Figure 6e), thus predicting a mechanism for transcriptional regulation of FgEC1 directly from promoter sequence.

## Discussion

Our new approach (RHIEPA) is able to generate regulatory maps for a variety of ascomycete fungi from genome sequences alone, without experimental data or whole genome alignments. For budding yeast, this allowed us to scale inferences of regulatory connections well-beyond the closely related *Saccharomyces sensu stricto* where genome alignments are available [98]. In this way, our approach is similar to phylogenetic footprinting [39], with the advantage that it develops a 256 dimensional motif-space that is easier to interpret than the 10,000s of k-mers used in systematic phylogenetic footprinting[39]. For relatively well-studied species, we took advantage of the large collections of known TF binding motifs to validate the learned motifs to assign TFs to them, and comparisons with TF overexpression experiments shows that the learned motif signals as effective (or more) as *in vivo* binding experiments at identifying responsive promoters (Supplementary Figure 1b). In some cases, we could understand the combinations of factors that control basic cellular processes, (such as the cell cycle and translation, Figure 3a) directly from the clustering of signals for motifs in promoter regions, and the clusters we obtain are comparable in their predictive power to machine learning classifiers trained on curated gene sets and genomics data (Supplementary Figure 2c).

Our unsupervised approach allows us to study regulatory function without relying on training data relating to biological function. Even with annotations or functional data, for pathways with small numbers of genes, such as glycolysis (24 genes annotated in *S. cerevisiae*) it would not be feasible to reliably learn the parameters of a motif of width 15 [5] or train a model to predict expression patterns from sequences in the genome[99]. By using orthologous non-coding sequences from more than 50 genomes to learn the motif-based representation, we augment the number of glycolysis promoters to more than 1000 [37]. By clustering and visualizing the representations, we can focus on strong patterns. Indeed, we identified a cluster of 16 promoters that contains 12 of the annotated glycolysis genes (Precision = 12/16 = 75% and Recall = 12/24 = 50%). Furthermore, these upstream regions contain signals for Gcr1, the known regulator of glycolysis as well as the upstream region for the *GCR1* gene, consistent with the known autoregulation[68].

Because RHIEPA uses genome sequences alone (no curated training data or genomics experiments are required) we can apply the clustering approach to poorly characterized species, including a plant pathogen difficult to study in the lab, where we predict both novel regulatory sequences and groups of potentially co-regulated genes, including uncharacterized genes that may function in novel pathways. For many of the promoter clusters in the RHIEPA representation, we find statistical enrichment of GO annotations comparable to classic gene expression analysis (Figures 3, 4 and 5) without any lab experiments. Again, because the analysis is based on sequence alone, we can identify pathways in relatively unstudied experimental conditions. This allowed us to identify putative pathways for parasex/meiosis in *C. albicans* (Figure 4), and mating and RNA-virus response in *N. crassa* (Figure 5). Thus, we believe that with sufficient evolutionary resolving power, unsupervised sequence-based analysis can represent a powerful alternative to direct functional genomics experiments for discovery in challenging organisms. Of course, where such data exists, future efforts can aim to integrate evolutionary conservation with experimental data[52,100,101].

An alternative approach to unsupervised learning from genome sequences is using large-language-models[40], one of which identified 51 motifs in *S. cerevisiae* 5’ regions through post-hoc motif-finding analysis, of which 18 were matched to known motifs [41]. Although 18 is far fewer than the 63 learned PWMs matched to known motifs here, it is not clear whether the difference is an advantage of a motif-based encoder relative to post-hoc motif-finding, or some other limitation of the large-language-model. More clear, however, is that the motif-based models here are trained on less than 300 million bases and have 80 thousand parameters, while the 5’ region large-language-model was trained on 13 billion bases and has 90 million parameters[41]. Thus, the computational expense of the motif-based approach is at least an order of magnitude less and the model complexity is 3 orders of magnitude less.

Despite the successful examples, matching learned motifs to databases has limitations. For example, the learned motif in the upstream regions of heat shock promoters (matrix 36 for *S. cerevisiae*, supplementary Figure 2, matrix 244 for *C. albicans,* Figure 4a), did not match any known motifs, even though they shows the well-known GAA repeats recognized by Hsf1[102]. Moreover, methionine pathway promoters in *S. cerevisiae* show a signal for matrix 233, whose best matches in JASPAR and cis-bp are Tye7 and Rtg3, respectively (E-value 4.83e-04 and 4.11e-03, respectively), TFs with similar specificity to one of the known regulators, Cbf1[103] (which is the second best match for matrix 233 in both databases, E-value 6.55e-03 and 1.96e-02, respectively). Thus, even interpretable motif-based approaches combined with extensive databases for well-studied organisms cannot definitively identify which transcription factor controls the pathways. Nevertheless, like high-throughput functional genomics experiments, the motif-based approach directly provides hypotheses that could be confirmed in simple experiments.

An oversimplification in our work (so far) is that we treat promoter and upstream regions interchangeably, taking the region upstream of the start codon to be the promoter, as has been done previously in fungal comparative genomics [39]. Even in fungal genomes, 5’ UTRs contain important post-transcriptional and translational regulatory elements[104] that are likely to be conserved and therefore provide signal for learning. However, because of the strong inductive bias of our encoder to learn transcription factor binding motifs (Figure 1b), we are not currently able to interpret these signals very well, although we do see motifs for the RNA-binding proteins PUF3 and PUF4[105] being learned in *S. cerevisiae*. RNA regulation is a possible explanation for some of the large clusters of upstream regions that show functional enrichments, but no strong signals for transcription factor motifs (e.g., Fusarium virulence and effector genes). Future applications of reverse homology could use more sophisticated encoders to model RNA structure and binding motifs[106]. For genomes with large non-coding regions, to apply a motif-based approach, the encoder will need to either operate over a large DNA region or identify the important regions via segmentation.

There are several important issues that arise when predicting regulatory networks from evolutionary comparisons of genome sequences. We rely on (i) a choice of species for comparison, (ii) evolutionary conservation of the regulatory network and (iii) ortholog assignments. Here we chose the species and number of orthologs arbitrarily based on the number of orthologs for each reference genome (see Methods). Connections in regulatory networks are well-known to turnover[16,74,75,77], such that similar regulation is achieved by different regulators in different lineages, and we noted three known examples in comparison of our *S. cerevisiae* and *C. albicans* representations (Glycolysis, GAL genes and ribosomal genes). Preliminary data indicates that such changes could be identified by averaging scores of known motifs over gene groups[31]. However, when we average the representations over the orthologs (Figure 1b) we are diluting out signals that are not conserved among all the species. Despite this, reverse homology and phylogenetic averaging appear to be robust to this issue [31,38], as long as a substantial fraction of species share the motif[31]. Of the issues listed above, for broad application to fungal genomes, we believe ortholog finding is currently the most pressing practical issue. Even within the groups of fungi considered here, current ortholog assignment software is difficult to scale to 100s or 1000s of genomes[107]. In our study, the ortholog finding was more computationally demanding than creating the RHIEPA representations (see Methods). It is already infeasible to apply high-confidence ortholog finding pipelines[107] to the 100,000s of bacterial genomes currently available. This is an advantage of unsupervised training of large-language-models[41] that do not rely on ortholog assignments.

Future motif-based representation-learning approaches that can leverage evolutionary conservation without requiring sequence alignment ***or*** ortholog assignments might be even more powerful tools for regulatory network inference.

## Methods

### Ortholog finding for fungal promoters

Y1000p [24] genomes were obtained from http://y1000plus.org. For filamentous fungi, 440 Sordariomycetes genomes were downloaded from NCBI using datasets tools (https://www.ncbi.nlm.nih.gov/datasets/). We ran OrthoFinder[107] using diamond_unltra_sens, --fewer-files and -p. Example codes for orthofinder runs are available at https://github.com/stajichlab/Y1000_orthologs.

We used NC12 as reference for *Neurospora* and ASM24013v3 as reference for *F. graminearum*. To choose the species to provide orthologs for each reference, we ranked species by how many reference genes had orthologs and then arbitrarily chose a number of species based on the coverage obtained. After ortholog finding, we noted redundant genome sequences and assemblies among the y1000+ genomes and manually removed them. The numbers were *S. cerevisiae*: 90 (1 redundant was filtered) species with at least 4,400 orthologs, *C. albicans*: 69 (7 redundant were filtered) species with at least 5,000 orthologs, *N. crassa*: 52 species with at least 6,500 orthologs, *F. graminearum*: 68 species with at least 8,000 orthologs (before filtering, see below). Ortholog coverage distributions per genome and per gene after selection of close genomes is shown in Supplementary Figure 3. The complete lists of species used are available as Supplementary File 2.

We extracted upstream sequence for each ortholog up to 800 bp or the next annotated coding sequence. For *F. graminearum*, we aligned the orthologs using mafft[108] we filtered out any ortholog sequences that were less than 0.01 subs. per site away from the reference, and using heuristics as for filtering protein orthologs[38].

Alignments of non-coding sequences in figures were done with mafft[108] and visualized using jalview[109].

### A motif-based encoder

Inspired by recent advances in fully interpretable encoders based on standard biophysical and bioinformatics models[32–35], we implemented standard bioinformatics motif-scanning steps in an end-to-end, trainable encoder using keras/tensorflow. Given an input of N orthologous sequences of varying lengths, we first pad shorter sequences to a fixed sequence length and then take the reverse complement. We scan both strands with the learned PWMs, reverse the order of scores of the reverse complement, and then take the better score of the two strands at each position. Although not indicated for simplicity in Figure 1b, in these steps we only consider matches in L – W positions, where W=15 is the motif width and, to avoid counting overlapping matches, we take the best match in a 10 nt window. We next scale the scores for each motif using a trainable scaling layer[35] and summarize the number and strength of matches using the trainable pooling layer[35]. This allows the model to choose the cutoff for each motif and decide if single strong matches or multiple weak matches should be considered for each motif. The outputs of these layers are weighted by an interaction matrix, scaled between 0 and 1[35]. We then applied batch normalization[110] which we found stabilized the training of the encoder.

To create a bona fide motif layer based on the standard 1d convolution layer, we implemented a constraint to ensure that a conv1d layer could be interpreted as a PWM. The kernel of a 1d convolution is simply a matrix of size W x C, where C are the channels (in our case C=4 for the 4 nucleotides of DNA represented as a matrix of indicator variables) and W is the width of the kernel (in our case the width of the PWM). Although a PWM can match the shape of a 1d convolution kernel (W x 4), the entries of a true PWM matrix, M, are defined as 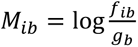 where *i* and *b* index the position and nucleotide, respectively, and *f* and *g* are probability distributions over nucleotides under the motif model and a background model, respectively[36]. Throughout we use *g* = (0.25, 0.25, 0.25, 0.25). To ensure that the probabilities under the motif model sum to 1 (and therefore each position has only 3 degrees of freedom, not 4), each position *i* of M should follow

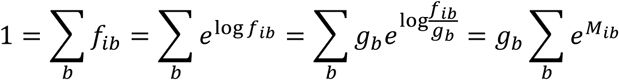

Where in the last step we took advantage of the assumption of equal background probabilities. Taking logs implies that for valid PWMs, log *g*_*b*_ + log ∑_*b*_ *e*^*Mib*^ = 0. We therefore implemented a PWM constraint in Keras, resetting the weights of the one-dimensional convolutional layer as follows:

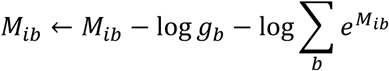

For example, consider the matrix of W=2, 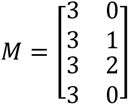. Neither of the columns is a valid position in a PWM (they contain log probability ratios greater than the background, but none smaller). Using base 2 logarithms for numerical convenience we have log_2_ *g*_*b*_ = −2, log_2_ ∑_*b*_ 2^*M*1*b*^ = 5 and log_2_ ∑_*b*_ 2^*M*2*b*^ = 3 for positions 1 and 2, respectively. Applying the PWM constraint, we have

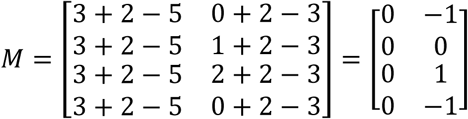

Converting these to motif probabilities using *f*_*ib*_ = *g*_*b*_2^*M_ib_*^ gives 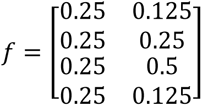 which are now valid probabilities. We added this PWM constraint to the convolutional layer in our model and confirmed empirically that our learned motifs are PWMs with *g* as defined.

### Learning non-coding sequence representations using reverse homology

Reverse homology is contrastive learning using evolution as the training signal[37,38]. It trains an encoder to tell apart members of a family of homologous (in this case orthologous, see above) sequences from randomly selected sequences. We largely followed the procedure suggested in previous work[38], with some minor simplifications. A single sequence is chosen as the ‘anchor’ (or positive) and it is encoded. n_f_ of the anchor’s homologs are sampled to represent the family, and their embeddings are simply per motif averaged to create the ‘family embedding’. Unlike previous work[38], we used family size n_f_ =4 and the same encoder for the family and the single sequences. We used each training sequence as the ‘anchor’ once in each epoch. During training, we filtered any sequences that were shorter than 10% of the max sequence length (L=800 bp) and removed any sequences that contain Ns (missing nucleotides). As in previous work[37], we use the infoNCE loss[111], which is the categorical cross entropy, *H*(*y, p*) = −∑*^B^_j_ y_j_* log *p_j_* where y is an indicator variable defined so that the positive anchor is 1 and all other single sequences are 0, and 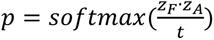 where · is the dot product of the family embedding *z_F_* and the single sequence embedding *z_F_*. In reverse homology, the infoNCE is using the dot product with the family embedding to classify the anchor as a homolog relative to all other anchors in the batch (which are negatives for that family). Throughout, we used temperature t=0.1 and batch size B= 256. We also applied reverse complement augmentation with probability 0.5, even though our encoder does consider both strands. With 256 motifs, the encoder includes 82688 parameters. We trained each model using Adam with learning rate 0.001 for a maximum of 100 epochs, and used a held-out validation set for early stopping (patience of 10 epochs). All models were trained on a workstation equipped with an Nvidia Quadro P6000. Example of a loss curve showing convergence can be found as Supplementary Figure 4.

Once the parameters have been trained, we encode promoter sequences into the 256-dimensional RHIEPA space using up to L=500bp upstream for each ortholog, as we found that this reduced the frequency of clustering together promoters of an adjacent genes (compared to using the full 800bp used in training the model.) We averaged the 256-dimensional representation over all available orthologs and filtered any upstream regions with less than 2 orthologs. Codes are available at https://github.com/alanmoses/promoter_reverse_homology

### Comparison of learned motifs to databases using TomTom

We converted the motifs to MEME format and uploaded them to the online MEME suite[44]. We searched JASPAR fungi [42]and cis-bp[43] for each species of interest. For the comparison of learned databases of motifs obtained using reverse homology (Figure 2b), we used E-value<1/256 to control for false matches. For the expert curated yetfasco PWMs [45], we used only the 244 high-confidence motifs, and E-value <1/244. TomTom outputs showing logos and E-values for all the motifs in each map are available as supplementary data.

For exploration of heatmaps, we assigned the learned motif to a transcription factor name from JASPAR fungi if the E-value was < 0.05. We did not use cis-bp for this as we found the names of motifs in that database were not usually interpretable.

### Comparison to TF overexpression of 15 activators

We downloaded binding evidence from Yeastract[47] and obtained transcription factor overexpression time courses [46]. For each of the 15 transcriptional activators, we identified any learned matrices in the S. cerevisiae RHIEPA representation that had that transcriptional activator as their best match using a TomTom [44] search (E-value <0.05) against JASPAR fungi[42]. We ranked the promoters by the scores for that motif. For the 5 of the 15 activators that had more than one best matching matrix, we ranked by the median of the scores (Rpn4 had 2, Sfp1 had 4, Ace2 had 2, Aft2 had 2 and Ste12 had 3) All the motifs for *S. cerevisiae* and their E-values from TomTom searches are available for download in html format at doi.org/10.5281/zenodo.14920043. For each transcriptional activator, we counted the number of genes that responded to overexpression (median log expression change over the time-course experiments >1) for the bound targets from Yeastract, or the same number of promoters ranked by their scores for the learned motif. To get an estimate of bona fide regulated promoters that don’t contain motifs, we counted the number of promoters bound by the TF and responding to that TF that were not in the top promoters ranked by motif scores. To test whether the low precision was due to the TF overexpression experiments missing most of the targets, we counted the number of responsive promoters among the top 50 promoters ranked by their motif scores.

### Systematic phylogenetic footprinting

We implemented a systematic phylogenetic footprinting strategy following RSAT tools[39] and applied it to our promoter ortholog sets for *S. cerevisiae*. For all 6-mers (and their reverse complements) including spacers up to 20 nucleotides (referred to as dyads[39]) that appear in the S. cerevisiae reference promoter (800 bp upstream of the start codon until the neighbouring gene), we counted the number of occurrences within the orthologous promoters and compared that to the expected proportion that dyad based on the counts in the entire dataset using the binomial distribution. Following[39] we converted the P-values to E-values using the number of dyads tested in that promoter and assigned the -log10 E-value to that promoter and that dyad if the E-value was less that 1, or assigned 0 otherwise. This procedure tested 41,600 dyads (and their reverse complements) for 5407 promoters. 41,547 dyads had at least one non-zero log10 E-value. This 41,600 dimensional representation for each promoter was reduced to 2 dimensions using UMAP and visualized (Supplementary Figure 1b) to confirm that it can identify groups of co-regulated promoters using genome sequences alone [39].

### Comparison of representations to gene expression data

We converted binding data (entire matrix of binding evidence downloaded from yeastract[47]) and netprophet3 regulatory network[54] to binary representation vectors where for each promoter where we have a 1 if a transcription factor binds it and 0 otherwise. We also tried using the confidence weights from netprophet3 as a high-dimensional representation, but obtained worse results. The phylogenetic footprinting representation was created as described above.

We used the sci-kit learn[112] neighbours module to identify nearest neighbours using cosine distance for each representation. We median centered the columns of each representation and the columns of the gene expression data[50] and confirmed that no expression experiments had >50% missing data. We set the remaining missing values to 0 (expression data is in log space) and confirmed that there were no experiments with 0 standard deviation.

For nearest neighbour expression experiment prediction, we replaced the observed gene expression values for a given experiment with the values of the nearest neighbour (or average of 50 nearest neighbours) in the representation space. We then computed the Pearson correlation between the nearest neighbour expression values and the observed gene expression values. This represents 1-nn regression with leave-one-out cross-validation.

For nearest neighbour prediction of co-expression, we computed the genes by genes correlation matrix and considered any gene pairs above correlation 0.5 to be co-expressed. Approximately 1% of gene pairs are considered co-expressed by this criterion. For each gene, we then checked whether a co-expressed gene was the nearest neighbour in the representation space (top 1 precision) or whether a co-expressed gene was among the 50 nearest neighbours (top 50 precision).

To obtain a baseline of sequence-based neighbours for this analysis, we ran an all by all BLAST search of the *S. cerevisiae* promoters with E-value cutoff = 10.0 and up to 5 matches to ensure that nearly all sequences had hits. We obtained the best hits for each promoter among the 5303 genes for which we had orthologs and gene expression data and assigned them as nearest neighbours. For the 25 promoters that did not have a BLAST hit (0.47%) we assigned a random nearest neighbour.

### Visualization and clustering of motif-based representations

We obtained gene descriptions from SGD[113] (budding yeast, *S. cerevisiae*) CGD[114] (*C. albicans*) Ensembl[79] (*N. crassa*, NC12, and *F. graminearum*, PH-1). We supplemented the *F. graminearum* annotations using annotations of our orthologs in neurospora.

We made UMAPs using the uwot package [115] in R and coloured points using grep of the gene names and descriptions. For example, for ‘proteasome’ in Figure 2s, we searched for any genes that had 20S proteasome or 26S proteasome in their gene description, while for ‘ribosome’, we searched for gene names that matched RPL or RPS. For PHO, ERG and MET pathways, we coloured genes with names matching PHO, ERG or MET, respectively.

For clustered heatmaps, we median centered and clustered using cluster3.0 (average linkage hierarchical clustering). Clustered representations were explored interactively in java treeview[116]. The datafiles for treeview for each of the 4 regulatory maps are provided as supplementary data.

### Comparison to cell cycle cluster to curated data and machine learning classifiers

We obtained a curated data set of 598 cell cycle regulated genes and 454 not cell cycle regulated genes[64] that were used for training machine learning classifiers based on selected *in vivo* transcription factor binding and promoter motifs and compared it to our cell cycle cluster, obtained by clustering the RHIEPA promoter representations (Figure 3, Supplementary File 1). We found that 289 of the cell cycle regulated genes fell into the cluster (True positive rate = 289/598 = 48%) and 18 of the genes in our cluster were among the 454 not cell cycle regulated (False positive rate = 18/454 = 4.0%). To compare these numbers to the performance reported for the machine learning classifiers[64], we divided their ROC curve using 20 x 20 equally spaced gridlines. The true positive rate and false positive rate for our cluster falls outside of the ROC curves they reported (Supplementary Figure 2c).

### Enrichment analyses

GO enrichment analysis was done using Gprofiler[56] using all annotated genes as the background for S. cerevisiae and all known genes in the species as the background for other species as they had much fewer annotations. Corrected P-values are reported in the text and gene lists are available in Supplementary File 1. Other data was tested for association using Fisher’s test in R[117]. Known targets of Neurospora transcription factors were obtained [83] and we used the low-throughput evidence as evidence of bona fide transcription factor-target interactions. Rhythmic promoters were identified using RNA pol binding and gene expression[84]. We parsed PHI-base[93] for *F. graminearum* genes that were annotated as ‘reduced virulence’: these are genes that result in reduced virulence when mutated. We obtained a list of *F. graminearum* genes that encode effectors from Fusarium Protein Toolkit (FPT)[94]. Gene lists for clusters discussed in the text are available as Supplementary File 1.

## Data and code availability

All data can be found at doi.org/10.5281/zenodo.14920043

Codes can be found at https://github.com/stajichlab/Y1000_orthologs and https://github.com/alanmoses/promoter_reverse_homology

## Acknowledgements

Research was performed using a GPU generously donated by Nvidia to AMM and hardware obtained through support to AMM from the Canada Foundation for Innovation. We thank members of the Moses lab, Knowles Lab, Philip Fradkin, Sanjana Bhatnagar and Dr. John Calarco for Discussions and Cameron Dufault for comments on the manuscript. We acknowledge Aqsa Alam for sharing data that was used for earlier versions of the analysis.

## Supplementary Figures

**Supplementary Figure 1.**
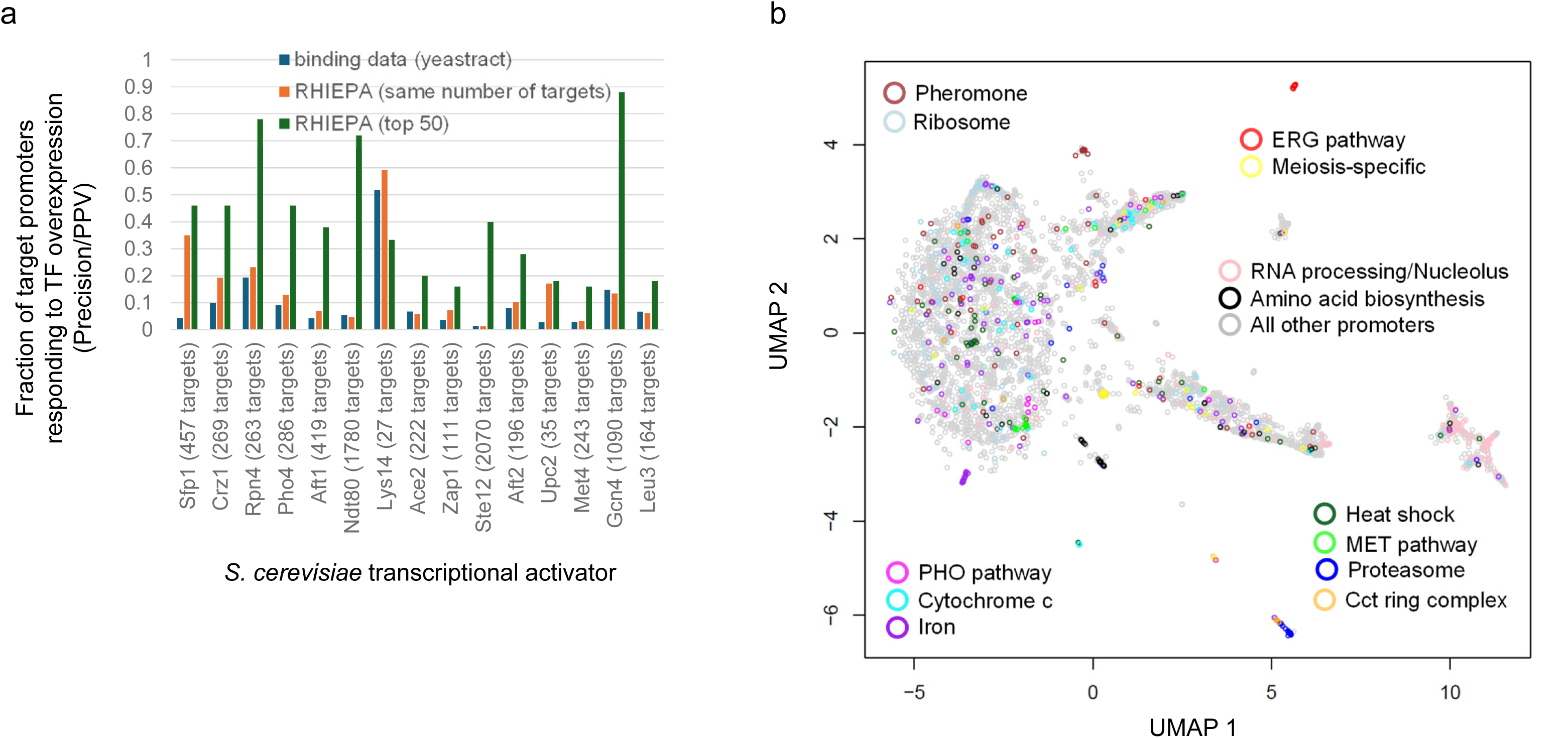
Other approaches to identify groups of co-regulated genes. (a) For 15 transcriptional activators from S. cerevisiae, the fraction of promoters that respond to TF overexpression is shown for in vivo bound targets from yeastract (blue bars), the same number of targets ranked by the median motif-based representation (RHIEPA) signals for those motifs identified in a TomTom search against JASPAR-fungi (orange bars) or the top 50 RHIEPA promoters (green bars). The number of in vivo bound targets for each transcriptional activator is indicated in the horizontal-axis label. (b) biological organization emerges in a 2D representation of the 41,600 dimensional phylogenetic footprinting space for *S. cerevisiae*. Each symbol represents a set or orthologous upstream regions. Colours indicate matches to the *S. cerevisiae* gene name or description.

**Supplementary Figure 2.**
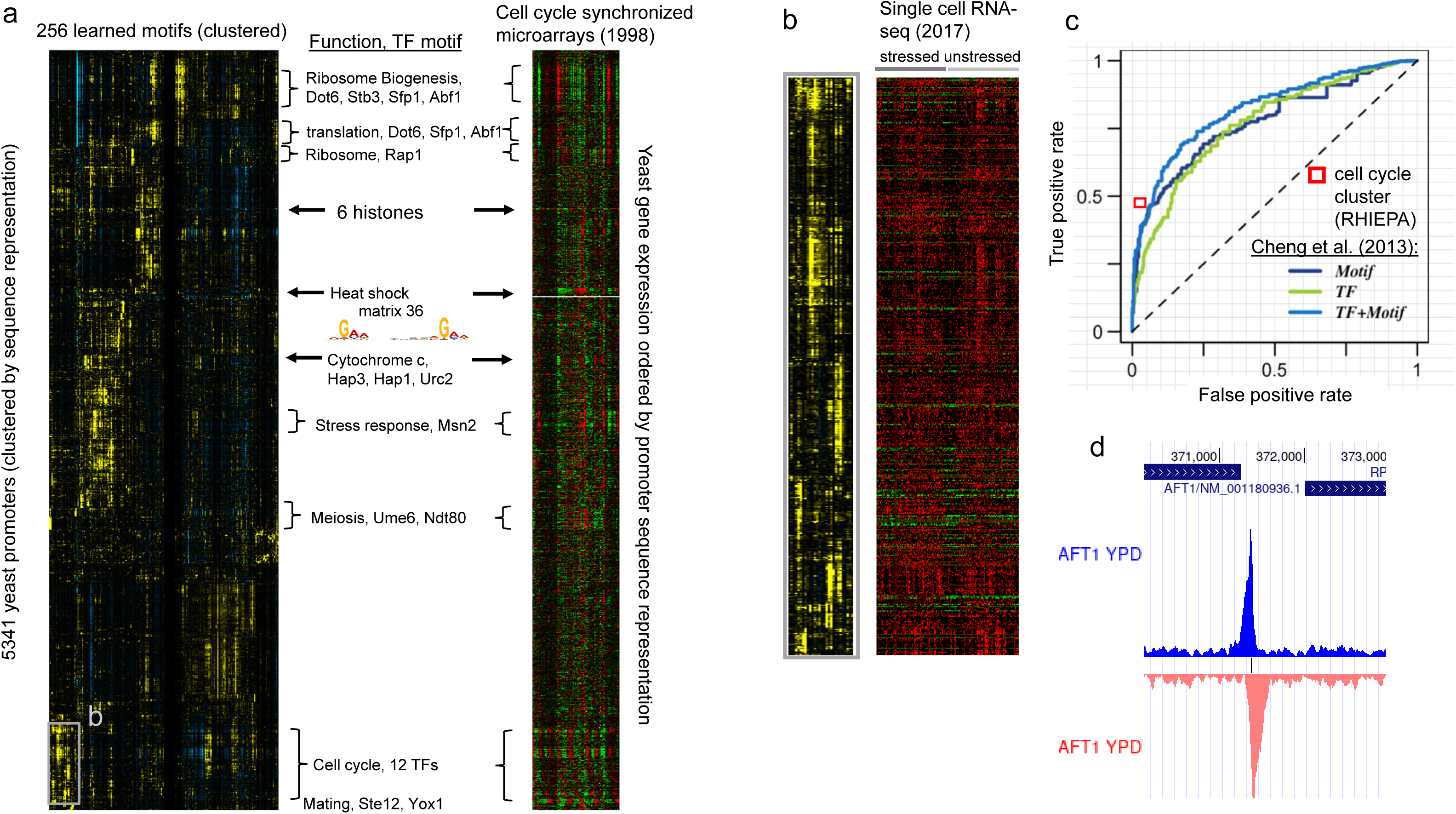
Additional evidence for understanding of the yeast regulatory code. a) comparison of the RHIEPA promoter representations to the entire cell synchronized microarray data. Gene expression data has been ordered according to the clustered RHIEPA promoter representation. In addition to the cell cycle cluster, clusters of promoters associated with translation, heat shock and general stress response, show changes in gene expression. These represent the various stress and growth responses due to the experimental conditions (alpha factor, temperature sensitive and other mutants) needed to synchronize the cells. Genes with meiosis regulator motifs in their promoters change expression in the Cdc28 mutant strains. a) genes with promoters in cell cycle clusters (Figure 3) show coordinated expression in single cell RNA-seq experiments. Rows (genes) in the single cell RNA-seq data have been ordered by the RHIEPA clustering of genes, and columns have been clustered based on the entire single-cell RNA-seq dataset. c) Equally spaced gridlines (grey lines) were superimposed on the original figure from Chen et al. 2013. Red box indicates the location of the cell cycle cluster with respect to the True positive rate and False positive rate using the training data from Cheng et al. 2013. The red box falls beyond the curves of all three of the machine learning classifiers they trained. d) Evidence of binding of Aft1 to the *AFT* promoter as displayed on the UCSC genome browser link from http://yeastepigenome.org/

**Supplementary Figure 3.**
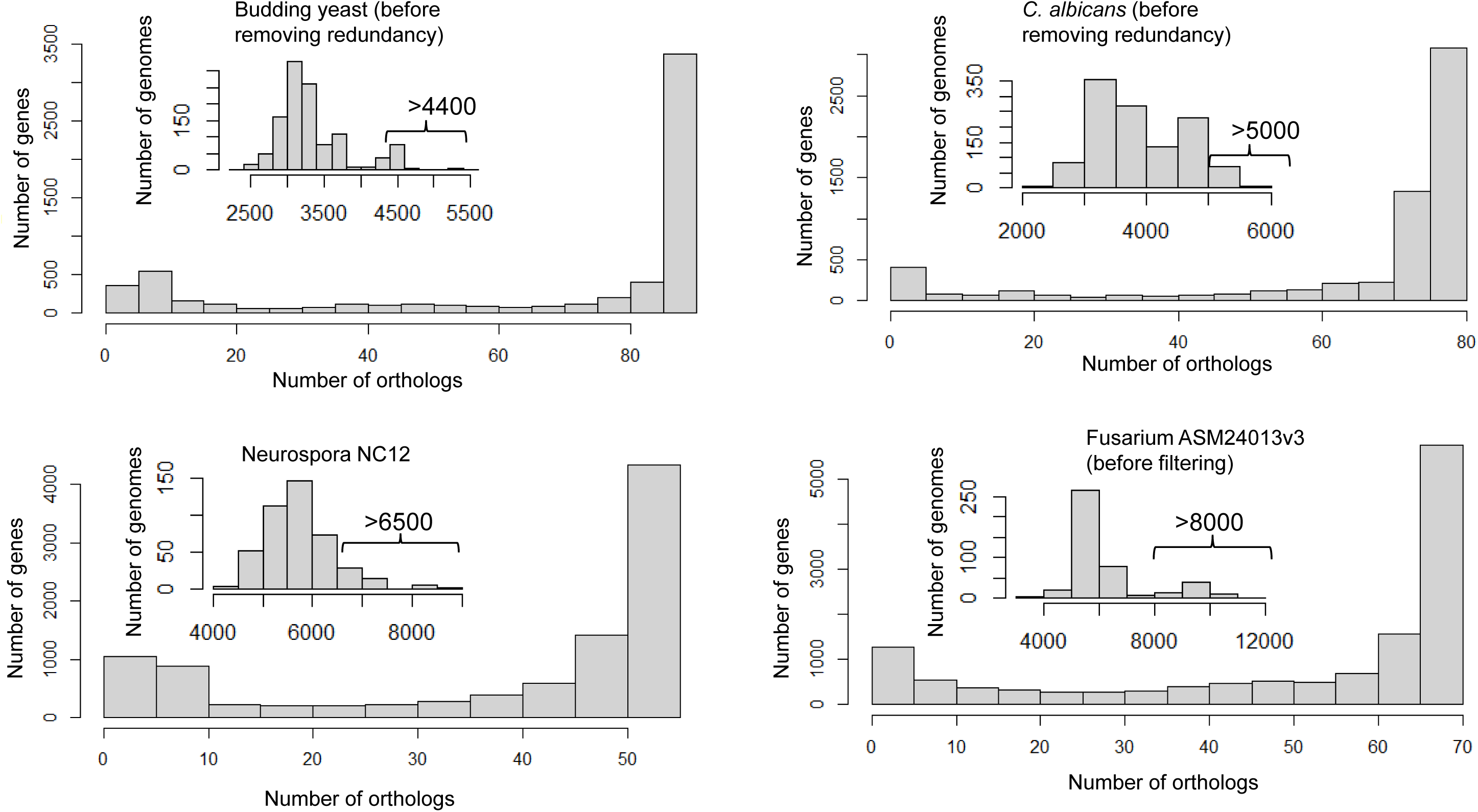
Ortholog coverage distribution. Histograms showing the distribution of the number of orthologs for each gene in the reference genomes among the selected closely related genomes. Inset in each panel is the overall distribution of number of orthologs per genome. Cutoff showing number of orthologs used to select closest genomes for further analysis is shown.

**Supplementary Figure 4.**
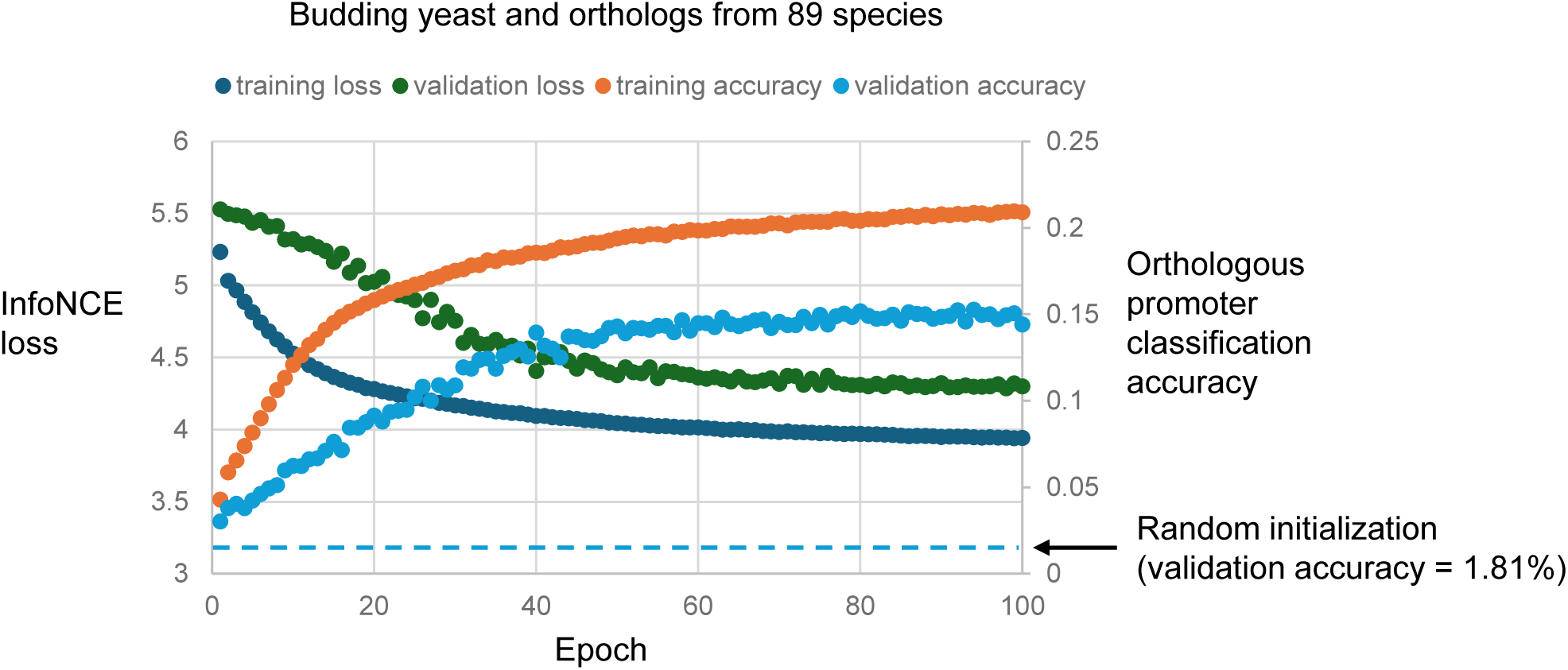
Example training curves. Example training information for the S. cerevisiae model. All 375 promoters (and orthologs) on chromosome 2 were held out as the validation sample. In this example, the model was trained for the full 100 epochs, as validation loss continued to decrease.

## Supplementary Files

**Supplementary File 1. Enrichment analysis.** This file is an Excel spreadsheet of gene lists and enrichment analysis for clusters discussed in the text. The first 4 tabs are the clusters discussed for each species. Each cluster is represented as two columns, gene IDs and gene descriptions. Subsequent columns include the Gprofiler results including the enriched annotation category, GO id, P-value, log P-value, number of genes in the genome with that annotation, number of genes in the cluster, number of genes in the cluster with that annotation and total number of genes. Below the fraction of genes in the cluster and genome are indicated, as well as their ratio, which is the fold enrichment used in the text. The final two tabs include the data used to compute enrichments for *F. graminearum* and *N. crassa* described in the text.

**Supplementary File 2. Species lists.** This file contains the names of the genome assemblies used for each of the 4 reference species.

## References

1. Aerts S. Computational strategies for the genome-wide identification of cis-regulatory elements and transcriptional targets. Curr Top Dev Biol. 2012;98: 121–145. doi:10.1016/B978-0-12-386499-4.00005-7

2. Kim S, Wysocka J. Deciphering the multi-scale, quantitative cis-regulatory code. Mol Cell. 2023; S1097–2765(22)01215–1. doi:10.1016/j.molcel.2022.12.032

3. Zeitlinger J. Seven myths of how transcription factors read the cis-regulatory code. Curr Opin Syst Biol. 2020;23: 22–31. doi:10.1016/j.coisb.2020.08.002

4. Wasserman WW, Sandelin A. Applied bioinformatics for the identification of regulatory elements. Nat Rev Genet. 2004;5: 276–287. doi:10.1038/nrg1315

5. Zia A, Moses AM. Towards a theoretical understanding of false positives in DNA motif finding. BMC Bioinformatics. 2012;13: 151. doi:10.1186/1471-2105-13-151

6. Moses AM, Chiang DY, Pollard DA, Iyer VN, Eisen MB. MONKEY: identifying conserved transcription-factor binding sites in multiple alignments using a binding site-specific evolutionary model. Genome Biol. 2004;5: R98. doi:10.1186/gb-2004-5-12-r98

7. Aerts S, Quan X-J, Claeys A, Sanchez MN, Tate P, Yan J, et al. Robust Target Gene Discovery through Transcriptome Perturbations and Genome-Wide Enhancer Predictions in Drosophila Uncovers a Regulatory Basis for Sensory Specification. PLOS Biol. 2010;8: e1000435. doi:10.1371/journal.pbio.1000435

8. Eddy SR. A Model of the Statistical Power of Comparative Genome Sequence Analysis. PLOS Biol. 2005;3: e10. doi:10.1371/journal.pbio.0030010

9. Kellis M, Patterson N, Endrizzi M, Birren B, Lander ES. Sequencing and comparison of yeast species to identify genes and regulatory elements. Nature. 2003;423: 241–254. doi:10.1038/nature01644

10. Xie X, Lu J, Kulbokas EJ, Golub TR, Mootha V, Lindblad-Toh K, et al. Systematic discovery of regulatory motifs in human promoters and 3’ UTRs by comparison of several mammals. Nature. 2005;434: 338–345. doi:10.1038/nature03441

11. Haudry A, Platts AE, Vello E, Hoen DR, Leclercq M, Williamson RJ, et al. An atlas of over 90,000 conserved noncoding sequences provides insight into crucifer regulatory regions. Nat Genet. 2013;45: 891–898. doi:10.1038/ng.2684

12. Kheradpour P, Stark A, Roy S, Kellis M. Reliable prediction of regulator targets using 12 Drosophila genomes. Genome Res. 2007;17: 1919–1931. doi:10.1101/gr.7090407

13. Ludwig MZ, Bergman C, Patel NH, Kreitman M. Evidence for stabilizing selection in a eukaryotic enhancer element. Nature. 2000;403: 564–567. doi:10.1038/35000615

14. Hare EE, Peterson BK, Iyer VN, Meier R, Eisen MB. Sepsid even-skipped enhancers are functionally conserved in Drosophila despite lack of sequence conservation. PLoS Genet. 2008;4: e1000106. doi:10.1371/journal.pgen.1000106

15. Doniger SW, Fay JC. Frequent gain and loss of functional transcription factor binding sites. PLoS Comput Biol. 2007;3: e99. doi:10.1371/journal.pcbi.0030099

16. Weirauch MT, Hughes TR. Conserved expression without conserved regulatory sequence: the more things change, the more they stay the same. Trends Genet TIG. 2010;26: 66–74. doi:10.1016/j.tig.2009.12.002

17. He X, Ling X, Sinha S. Alignment and Prediction of cis-Regulatory Modules Based on a Probabilistic Model of Evolution. PLOS Comput Biol. 2009;5: e1000299. doi:10.1371/journal.pcbi.1000299

18. Lunter G, Rocco A, Mimouni N, Heger A, Caldeira A, Hein J. Uncertainty in homology inferences: assessing and improving genomic sequence alignment. Genome Res. 2008;18: 298–309. doi:10.1101/gr.6725608

19. Ray P, Shringarpure S, Kolar M, Xing EP. CSMET: Comparative Genomic Motif Detection via Multi-Resolution Phylogenetic Shadowing. PLOS Comput Biol. 2008;4: e1000090. doi:10.1371/journal.pcbi.1000090

20. Arnold P, Erb I, Pachkov M, Molina N, van Nimwegen E. MotEvo: integrated Bayesian probabilistic methods for inferring regulatory sites and motifs on multiple alignments of DNA sequences. Bioinforma Oxf Engl. 2012;28: 487–494. doi:10.1093/bioinformatics/btr695

21. Glenwinkel L, Wu D, Minevich G, Hobert O. TargetOrtho: a phylogenetic footprinting tool to identify transcription factor targets. Genetics. 2014;197: 61–76. doi:10.1534/genetics.113.160721

22. Gordân R, Narlikar L, Hartemink AJ. Finding regulatory DNA motifs using alignment-free evolutionary conservation information. Nucleic Acids Res. 2010;38: e90. doi:10.1093/nar/gkp1166

23. Christmas MJ, Kaplow IM, Genereux DP, Dong MX, Hughes GM, Li X, et al. Evolutionary constraint and innovation across hundreds of placental mammals. Science. 2023;380: eabn3943. doi:10.1126/science.abn3943

24. Opulente DA, LaBella AL, Harrison M-C, Wolters JF, Liu C, Li Y, et al. Genomic factors shape carbon and nitrogen metabolic niche breadth across Saccharomycotina yeasts. Science. 2024;384: eadj4503. doi:10.1126/science.adj4503

25. Kim BY, Wang JR, Miller DE, Barmina O, Delaney E, Thompson A, et al. Highly contiguous assemblies of 101 drosophilid genomes. Coop G, Wittkopp PJ, Sackton TB, editors. eLife. 2021;10: e66405. doi:10.7554/eLife.66405

26. Hahn S, Young ET. Transcriptional Regulation in Saccharomyces cerevisiae: Transcription Factor Regulation and Function, Mechanisms of Initiation, and Roles of Activators and Coactivators. Genetics. 2011;189: 705–736. doi:10.1534/genetics.111.127019

27. Rodriguez DL, Quail MM, Hernday AD, Nobile CJ. Transcriptional Circuits Regulating Developmental Processes in Candida albicans. Front Cell Infect Microbiol. 2020;10: 605711. doi:10.3389/fcimb.2020.605711

28. Gostinčar C, Stajich JE, Gunde-Cimerman N. Extremophilic and extremotolerant fungi. Curr Biol. 2023;33: R752–R756. doi:10.1016/j.cub.2023.06.011

29. Case NT, Berman J, Blehert DS, Cramer RA, Cuomo C, Currie CR, et al. The future of fungi: threats and opportunities. G3 Bethesda Md. 2022;12: jkac224. doi:10.1093/g3journal/jkac224

30. Koo PK, Eddy SR. Representation learning of genomic sequence motifs with convolutional neural networks. PLOS Comput Biol. 2019;15: e1007560. doi:10.1371/journal.pcbi.1007560

31. Alam A, Duncan AG, Mitchell JA, Moses AM. Functional similarity of non-coding regions is revealed in phylogenetic average motif score representations. bioRxiv; 2023. p. 2023.04.09.536185. doi:10.1101/2023.04.09.536185

32. Tareen A, Kinney JB. Biophysical models of cis-regulation as interpretable neural networks. arXiv; 2020. doi:10.48550/arXiv.2001.03560

33. de Boer CG, Vaishnav ED, Sadeh R, Abeyta EL, Friedman N, Regev A. Deciphering eukaryotic gene-regulatory logic with 100 million random promoters. Nat Biotechnol. 2020;38: 56–65. doi:10.1038/s41587-019-0315-8

34. Novakovsky G, Fornes O, Saraswat M, Mostafavi S, Wasserman WW. ExplaiNN: interpretable and transparent neural networks for genomics. bioRxiv; 2022. p. 2022.05.20.492818. doi:10.1101/2022.05.20.492818

35. Balcı AT, Ebeid MM, Benos PV, Kostka D, Chikina M. An intrinsically interpretable neural network architecture for sequence to function learning. bioRxiv; 2023. p. 2023.01.25.525572. doi:10.1101/2023.01.25.525572

36. Stormo GD. DNA binding sites: representation and discovery. Bioinformatics. 2000;16: 16–23. doi:10.1093/bioinformatics/16.1.16

37. Lu AX, Lu AX, Moses A. Evolution Is All You Need: Phylogenetic Augmentation for Contrastive Learning. arXiv; 2020. doi:10.48550/arXiv.2012.13475

38. Lu AX, Lu AX, Pritišanac I, Zarin T, Forman-Kay JD, Moses AM. Discovering molecular features of intrinsically disordered regions by using evolution for contrastive learning. PLOS Comput Biol. 2022;18: e1010238. doi:10.1371/journal.pcbi.1010238

39. Brohée S, Janky R, Abdel-Sater F, Vanderstocken G, André B, van Helden J. Unraveling networks of co-regulated genes on the sole basis of genome sequences. Nucleic Acids Res. 2011;39: 6340–6358. doi:10.1093/nar/gkr264

40. Consens ME, Dufault C, Wainberg M, Forster D, Karimzadeh M, Goodarzi H, et al. To Transformers and Beyond: Large Language Models for the Genome. arXiv; 2023. doi:10.48550/arXiv.2311.07621

41. Karollus A, Hingerl J, Gankin D, Grosshauser M, Klemon K, Gagneur J. Species-aware DNA language models capture regulatory elements and their evolution. Genome Biol. 2024;25: 83. doi:10.1186/s13059-024-03221-x

42. Castro-Mondragon JA, Riudavets-Puig R, Rauluseviciute I, Lemma RB, Turchi L, Blanc-Mathieu R, et al. JASPAR 2022: the 9th release of the open-access database of transcription factor binding profiles. Nucleic Acids Res. 2022;50: D165–D173. doi:10.1093/nar/gkab1113

43. Lambert SA, Yang AWH, Sasse A, Cowley G, Albu M, Caddick MX, et al. Similarity regression predicts evolution of transcription factor sequence specificity. Nat Genet. 2019;51: 981–989. doi:10.1038/s41588-019-0411-1

44. Bailey TL, Boden M, Buske FA, Frith M, Grant CE, Clementi L, et al. MEME SUITE: tools for motif discovery and searching. Nucleic Acids Res. 2009;37: W202–208. doi:10.1093/nar/gkp335

45. de Boer CG, Hughes TR. YeTFaSCo: a database of evaluated yeast transcription factor sequence specificities. Nucleic Acids Res. 2012;40: D169–179. doi:10.1093/nar/gkr993

46. Hackett SR, Baltz EA, Coram M, Wranik BJ, Kim G, Baker A, et al. Learning causal networks using inducible transcription factors and transcriptome-wide time series. Mol Syst Biol. 2020;16: e9174. doi:10.15252/msb.20199174

47. Teixeira MC, Viana R, Palma M, Oliveira J, Galocha M, Mota MN, et al. YEASTRACT+: a portal for the exploitation of global transcription regulation and metabolic model data in yeast biotechnology and pathogenesis. Nucleic Acids Res. 2023;51: D785–D791. doi:10.1093/nar/gkac1041

48. Mahendrawada L, Warfield L, Donczew R, Hahn S. Low overlap of transcription factor DNA binding and regulatory targets. Nature. 2025;642: 796–804. doi:10.1038/s41586-025-08916-0

49. Gonsalves SE, Moses AM, Razak Z, Robert F, Westwood JT. Whole-genome analysis reveals that active heat shock factor binding sites are mostly associated with non-heat shock genes in Drosophila melanogaster. PloS One. 2011;6: e15934. doi:10.1371/journal.pone.0015934

50. Kemmeren P, Sameith K, van de Pasch LAL, Benschop JJ, Lenstra TL, Margaritis T, et al. Large-scale genetic perturbations reveal regulatory networks and an abundance of gene-specific repressors. Cell. 2014;157: 740–752. doi:10.1016/j.cell.2014.02.054

51. Karollus A, Mauermeier T, Gagneur J. Current sequence-based models capture gene expression determinants in promoters but mostly ignore distal enhancers. Genome Biol. 2023;24: 56. doi:10.1186/s13059-023-02899-9

52. Zhang S, Pyne S, Pietrzak S, Halberg S, McCalla SG, Siahpirani AF, et al. Inference of cell type-specific gene regulatory networks on cell lineages from single cell omic datasets. Nat Commun. 2023;14: 3064. doi:10.1038/s41467-023-38637-9

53. Skok Gibbs C, Jackson CA, Saldi G-A, Tjärnberg A, Shah A, Watters A, et al. High-performance single-cell gene regulatory network inference at scale: the Inferelator 3.0. Bioinformatics. 2022;38: 2519–2528. doi:10.1093/bioinformatics/btac117

54. Abid D, Brent MR. NetProphet 3: a machine learning framework for transcription factor network mapping and multi-omics integration. Bioinformatics. 2023;39: btad038. doi:10.1093/bioinformatics/btad038

55. Eisen MB, Spellman PT, Brown PO, Botstein D. Cluster analysis and display of genome-wide expression patterns. Proc Natl Acad Sci U S A. 1998;95: 14863–14868. doi:10.1073/pnas.95.25.14863

56. Kolberg L, Raudvere U, Kuzmin I, Adler P, Vilo J, Peterson H. g:Profiler-interoperable web service for functional enrichment analysis and gene identifier mapping (2023 update). Nucleic Acids Res. 2023;51: W207–W212. doi:10.1093/nar/gkad347

57. Shore D, Zencir S, Albert B. Transcriptional control of ribosome biogenesis in yeast: links to growth and stress signals. Biochem Soc Trans. 2021;49: 1589–1599. doi:10.1042/BST20201136

58. Hinnebusch AG, Natarajan K. Gcn4p, a master regulator of gene expression, is controlled at multiple levels by diverse signals of starvation and stress. Eukaryot Cell. 2002;1: 22–32. doi:10.1128/EC.01.1.22-32.2002

59. McFarland MR, Keller CD, Childers BM, Adeniyi SA, Corrigall H, Raguin A, et al. The molecular aetiology of tRNA synthetase depletion: induction of a GCN4 amino acid starvation response despite homeostatic maintenance of charged tRNA levels. Nucleic Acids Res. 2020;48: 3071–3088. doi:10.1093/nar/gkaa055

60. Spellman PT, Sherlock G, Zhang MQ, Iyer VR, Anders K, Eisen MB, et al. Comprehensive identification of cell cycle-regulated genes of the yeast Saccharomyces cerevisiae by microarray hybridization. Mol Biol Cell. 1998;9: 3273–3297. doi:10.1091/mbc.9.12.3273

61. Gasch AP, Yu FB, Hose J, Escalante LE, Place M, Bacher R, et al. Single-cell RNA sequencing reveals intrinsic and extrinsic regulatory heterogeneity in yeast responding to stress. PLoS Biol. 2017;15: e2004050. doi:10.1371/journal.pbio.2004050

62. Koch C, Moll T, Neuberg M, Ahorn H, Nasmyth K. A role for the transcription factors Mbp1 and Swi4 in progression from G1 to S phase. Science. 1993;261: 1551–1557. doi:10.1126/science.8372350

63. Pramila T, Wu W, Miles S, Noble WS, Breeden LL. The Forkhead transcription factor Hcm1 regulates chromosome segregation genes and fills the S-phase gap in the transcriptional circuitry of the cell cycle. Genes Dev. 2006;20: 2266–2278. doi:10.1101/gad.1450606

64. Cheng C, Fu Y, Shen L, Gerstein M. Identification of yeast cell cycle regulated genes based on genomic features. BMC Syst Biol. 2013;7: 70. doi:10.1186/1752-0509-7-70

65. Horak CE, Luscombe NM, Qian J, Bertone P, Piccirrillo S, Gerstein M, et al. Complex transcriptional circuitry at the G1/S transition in Saccharomyces cerevisiae. Genes Dev. 2002;16: 3017–3033. doi:10.1101/gad.1039602

66. Ramos-Alonso L, Romero AM, Martínez-Pastor MT, Puig S. Iron Regulatory Mechanisms in Saccharomyces cerevisiae. Front Microbiol. 2020;11: 582830. doi:10.3389/fmicb.2020.582830

67. Rossi MJ, Kuntala PK, Lai WKM, Yamada N, Badjatia N, Mittal C, et al. A high-resolution protein architecture of the budding yeast genome. Nature. 2021;592: 309–314. doi:10.1038/s41586-021-03314-8

68. Sasaki H, Kishimoto T, Mizuno T, Shinzato T, Uemura H. Expression of GCR1, the transcriptional activator of glycolytic enzyme genes in the yeast Saccharomyces cerevisiae, is positively autoregulated by Gcr1p. Yeast Chichester Engl. 2005;22: 305–319. doi:10.1002/yea.1212

69. Abramova NE, Cohen BD, Sertil O, Kapoor R, Davies KJ, Lowry CV. Regulatory mechanisms controlling expression of the DAN/TIR mannoprotein genes during anaerobic remodeling of the cell wall in Saccharomyces cerevisiae. Genetics. 2001;157: 1169–1177. doi:10.1093/genetics/157.3.1169

70. Prugar E, Burnett C, Chen X, Hollingsworth NM. Coordination of Double Strand Break Repair and Meiotic Progression in Yeast by a Mek1-Ndt80 Negative Feedback Loop. Genetics. 2017;206: 497–512. doi:10.1534/genetics.117.199703

71. Zhao H, Eide DJ. Zap1p, a metalloregulatory protein involved in zinc-responsive transcriptional regulation in Saccharomyces cerevisiae. Mol Cell Biol. 1997;17: 5044–5052. doi:10.1128/MCB.17.9.5044

72. Le Crom S, Devaux F, Marc P, Zhang X, Moye-Rowley WS, Jacq C. New insights into the pleiotropic drug resistance network from genome-wide characterization of the YRR1 transcription factor regulation system. Mol Cell Biol. 2002;22: 2642–2649. doi:10.1128/MCB.22.8.2642-2649.2002

73. Askew C, Sellam A, Epp E, Hogues H, Mullick A, Nantel A, et al. Transcriptional regulation of carbohydrate metabolism in the human pathogen Candida albicans. PLoS Pathog. 2009;5: e1000612. doi:10.1371/journal.ppat.1000612

74. Hogues H, Lavoie H, Sellam A, Mangos M, Roemer T, Purisima E, et al. Transcription factor substitution during the evolution of fungal ribosome regulation. Mol Cell. 2008;29: 552–562. doi:10.1016/j.molcel.2008.02.006

75. Lavoie H, Hogues H, Mallick J, Sellam A, Nantel A, Whiteway M. Evolutionary tinkering with conserved components of a transcriptional regulatory network. PLoS Biol. 2010;8: e1000329. doi:10.1371/journal.pbio.1000329

76. Sun X, Yu J, Zhu C, Mo X, Sun Q, Yang D, et al. Recognition of galactose by a scaffold protein recruits a transcriptional activator for the GAL regulon induction in Candida albicans. Rokas A, Struhl K, Whiteway M, editors. eLife. 2023;12: e84155. doi:10.7554/eLife.84155

77. Martchenko M, Levitin A, Hogues H, Nantel A, Whiteway M. Transcriptional Rewiring of Fungal Galactose-Metabolism Circuitry. Curr Biol. 2007;17: 1007–1013. doi:10.1016/j.cub.2007.05.017

78. Bennett RJ. The parasexual lifestyle of Candida albicans. Curr Opin Microbiol. 2015;28: 10–17. doi:10.1016/j.mib.2015.06.017

79. Ensembl Fungi. [cited 24 Feb 2025]. Available: https://fungi.ensembl.org/index.html

80. Gasch AP, Moses AM, Chiang DY, Fraser HB, Berardini M, Eisen MB. Conservation and evolution of cis-regulatory systems in ascomycete fungi. PLoS Biol. 2004;2: e398. doi:10.1371/journal.pbio.0020398

81. Hynes MJ, Murray SL, Duncan A, Khew GS, Davis MA. Regulatory genes controlling fatty acid catabolism and peroxisomal functions in the filamentous fungus Aspergillus nidulans. Eukaryot Cell. 2006;5: 794–805. doi:10.1128/EC.5.5.794-805.2006

82. Lamb TM, Finch KE, Bell-Pedersen D. The *Neurospora crassa* OS MAPK pathway-activated transcription factor ASL-1 contributes to circadian rhythms in pathway responsive clock-controlled genes. Fungal Genet Biol. 2012;49: 180–188. doi:10.1016/j.fgb.2011.12.006

83. Hu Y, Qin Y, Liu G. Collection and Curation of Transcriptional Regulatory Interactions in Aspergillus nidulans and Neurospora crassa Reveal Structural and Evolutionary Features of the Regulatory Networks. Front Microbiol. 2018;9: 27. doi:10.3389/fmicb.2018.00027

84. Sancar C, Sancar G, Ha N, Cesbron F, Brunner M. Dawn- and dusk-phased circadian transcription rhythms coordinate anabolic and catabolic functions in Neurospora. BMC Biol. 2015;13: 17. doi:10.1186/s12915-015-0126-4

85. Wu VW, Thieme N, Huberman LB, Dietschmann A, Kowbel DJ, Lee J, et al. The regulatory and transcriptional landscape associated with carbon utilization in a filamentous fungus. Proc Natl Acad Sci U S A. 2020;117: 6003–6013. doi:10.1073/pnas.1915611117

86. Philley ML, Staben C. Functional analyses of the Neurospora crassa MT a-1 mating type polypeptide. Genetics. 1994;137: 715–722. doi:10.1093/genetics/137.3.715

87. Shiu PK, Raju NB, Zickler D, Metzenberg RL. Meiotic silencing by unpaired DNA. Cell. 2001;107: 905–916. doi:10.1016/s0092-8674(01)00609-2

88. Shiu PKT, Zickler D, Raju NB, Ruprich-Robert G, Metzenberg RL. SAD-2 is required for meiotic silencing by unpaired DNA and perinuclear localization of SAD-1 RNA-directed RNA polymerase. Proc Natl Acad Sci U S A. 2006;103: 2243–2248. doi:10.1073/pnas.0508896103

89. Lee DW, Pratt RJ, McLaughlin M, Aramayo R. An Argonaute-Like Protein Is Required for Meiotic Silencing. Genetics. 2003;164: 821–828. doi:10.1093/genetics/164.2.821

90. Wang Z, Lopez-Giraldez F, Slot J, Yarden O, Trail F, Townsend JP. Secondary Metabolism Gene Clusters Exhibit Increasingly Dynamic and Differential Expression during Asexual Growth, Conidiation, and Sexual Development in Neurospora crassa. mSystems. 2022;7: e0023222. doi:10.1128/msystems.00232-22

91. Honda S, Eusebio-Cope A, Miyashita S, Yokoyama A, Aulia A, Shahi S, et al. Establishment of Neurospora crassa as a model organism for fungal virology. Nat Commun. 2020;11: 5627. doi:10.1038/s41467-020-19355-y

92. Alouane T, Rimbert H, Bormann J, González-Montiel GA, Loesgen S, Schäfer W, et al. Comparative Genomics of Eight Fusarium graminearum Strains with Contrasting Aggressiveness Reveals an Expanded Open Pangenome and Extended Effector Content Signatures. Int J Mol Sci. 2021;22: 6257. doi:10.3390/ijms22126257

93. Urban M, Cuzick A, Seager J, Nonavinakere N, Sahoo J, Sahu P, et al. PHI-base - the multi-species pathogen-host interaction database in 2025. Nucleic Acids Res. 2025;53: D826–D838. doi:10.1093/nar/gkae1084

94. Kim H-S, Haley OC, Portwood Ii JL, Harding S, Proctor RH, Woodhouse MR, et al. Fusarium Protein Toolkit: a web-based resource for structural and variant analysis of Fusarium species. BMC Microbiol. 2024;24: 326. doi:10.1186/s12866-024-03480-5

95. Cai L, Xu X, Dong Y, Jin Y, Rashad YM, Ma D, et al. Roles of Three FgPel Genes in the Development and Pathogenicity Regulation of Fusarium graminearum. J Fungi. 2024;10: 666. doi:10.3390/jof10100666

96. Gu Q, Zhang C, Liu X, Ma Z. A transcription factor FgSte12 is required for pathogenicity in Fusarium graminearum. Mol Plant Pathol. 2015;16: 1–13. doi:10.1111/mpp.12155

97. Shang S, He Y, Hu Q, Fang Y, Cheng S, Zhang C-J. Fusarium graminearum effector FgEC1 targets wheat TaGF14b protein to suppress TaRBOHD-mediated ROS production and promote infection. J Integr Plant Biol. 2024;66: 2288–2303. doi:10.1111/jipb.13752

98. Scannell DR, Zill OA, Rokas A, Payen C, Dunham MJ, Eisen MB, et al. The Awesome Power of Yeast Evolutionary Genetics: New Genome Sequences and Strain Resources for the Saccharomyces sensu stricto Genus. G3 Bethesda Md. 2011;1: 11–25. doi:10.1534/g3.111.000273

99. de Boer CG, Taipale J. Hold out the genome: a roadmap to solving the cis-regulatory code. Nature. 2024;625: 41–50. doi:10.1038/s41586-023-06661-w

100. Carriel CC, Pyne S, Halberg-Spencer SA, Park SC, Seo H, Schmidt A, et al. A network-based model of Aspergillus fumigatus elucidates regulators of development and defensive natural products of an opportunistic pathogen. bioRxiv. 2023; 2023–05.

101. Zhang S, Knaack S, Roy S. Enabling Studies of Genome-Scale Regulatory Network Evolution in Large Phylogenies with MRTLE. Methods Mol Biol Clifton NJ. 2022;2477: 439–455. doi:10.1007/978-1-0716-2257-5_24

102. Sakurai H, Takemori Y. Interaction between Heat Shock Transcription Factors (HSFs) and Divergent Binding Sequences: BINDING SPECIFICITIES OF YEAST HSFs AND HUMAN HSF1*. J Biol Chem. 2007;282: 13334–13341. doi:10.1074/jbc.M611801200

103. Petti AA, McIsaac RS, Ho-Shing O, Bussemaker HJ, Botstein D. Combinatorial control of diverse metabolic and physiological functions by transcriptional regulators of the yeast sulfur assimilation pathway. Mol Biol Cell. 2012;23: 3008–3024. doi:10.1091/mbc.E12-03-0233

104. Dvir S, Velten L, Sharon E, Zeevi D, Carey LB, Weinberger A, et al. Deciphering the rules by which 5’-UTR sequences affect protein expression in yeast. Proc Natl Acad Sci U S A. 2013;110: E2792–2801. doi:10.1073/pnas.1222534110

105. Gerber AP, Herschlag D, Brown PO. Extensive association of functionally and cytotopically related mRNAs with Puf family RNA-binding proteins in yeast. PLoS Biol. 2004;2: E79. doi:10.1371/journal.pbio.0020079

106. Saman Booy M, Ilin A, Orponen P. RNA secondary structure prediction with convolutional neural networks. BMC Bioinformatics. 2022;23: 58. doi:10.1186/s12859-021-04540-7

107. Emms DM, Kelly S. OrthoFinder: phylogenetic orthology inference for comparative genomics. Genome Biol. 2019;20: 238. doi:10.1186/s13059-019-1832-y

108. Katoh K, Standley DM. MAFFT multiple sequence alignment software version 7: improvements in performance and usability. Mol Biol Evol. 2013;30: 772–780. doi:10.1093/molbev/mst010

109. Waterhouse AM, Procter JB, Martin DMA, Clamp M, Barton GJ. Jalview Version 2--a multiple sequence alignment editor and analysis workbench. Bioinforma Oxf Engl. 2009;25: 1189–1191. doi:10.1093/bioinformatics/btp033

110. Ioffe S, Szegedy C. Batch Normalization: Accelerating Deep Network Training by Reducing Internal Covariate Shift. arXiv; 2015. doi:10.48550/arXiv.1502.03167

111. Oord A van den, Li Y, Vinyals O. Representation Learning with Contrastive Predictive Coding. arXiv; 2019. doi:10.48550/arXiv.1807.03748

112. Pedregosa F, Varoquaux G, Gramfort A, Michel V, Thirion B, Grisel O, et al. Scikit-learn: Machine Learning in Python. J Mach Learn Res. 2011;12: 2825–2830.

113. Engel SR, Aleksander S, Nash RS, Wong ED, Weng S, Miyasato SR, et al. Saccharomyces Genome Database: Advances in Genome Annotation, Expanded Biochemical Pathways, and Other Key Enhancements. Genetics. 2024; iyae185. doi:10.1093/genetics/iyae185

114. Skrzypek MS, Binkley J, Binkley G, Miyasato SR, Simison M, Sherlock G. The Candida Genome Database (CGD): incorporation of Assembly 22, systematic identifiers and visualization of high throughput sequencing data. Nucleic Acids Res. 2017;45: D592– D596. doi:10.1093/nar/gkw924

115. Melville J. jlmelville/uwot. 2025. Available: https://github.com/jlmelville/uwot

116. Saldanha AJ. Java Treeview—extensible visualization of microarray data. Bioinformatics. 2004;20: 3246–3248. doi:10.1093/bioinformatics/bth349

117. Ihaka R, Gentleman R. R: A Language for Data Analysis and Graphics. J Comput Graph Stat. 1996;5: 299–314. doi:10.2307/1390807

